# Restriction of access to the central cavity is a major contributor to substrate selectivity in plant ABCG transporters

**DOI:** 10.1101/2022.08.16.503940

**Authors:** Konrad Pakuła, Carlos Sequeiros-Borja, Wanda Biała-Leonhard, Aleksandra Pawela, Joanna Banasiak, Aurélien Bailly, Marcin Radom, Markus Geisler, Jan Brezovsky, Michał Jasiński

**Author notes:** These authors contributed equally to the work (Konrad Pakuła, Carlos Sequeiros-Borja, Wanda Biała-Leonhard). For correspondence: **Michał Jasiński**, Department of Plant Molecular Physiology, Institute of Bioorganic Chemistry, Polish Academy of Sciences, Z. Noskowskiego 12/14, 61-704 Poznan, Poland; **Jan Brezovsky**, Laboratory of Biomolecular Interactions and Transport, Department of Gene Expression, Institute of Molecular Biology and Biotechnology, Faculty of Biology, Adam Mickiewicz University, Uniwersytetu Poznanskiego 6, 61-614 Poznan, Poland, and International Institute of Molecular and Cell Biology in Warsaw, 02-109 Warsaw, Poland).

## Abstract

ABCG46 of the legume *Medicago truncatula* is an ABC-type transporter responsible for highly selective translocation of the phenylpropanoids, 4-coumarate and liquiritigenin, over the plasma membrane. To investigate molecular determinants of the observed substrate selectivity, we applied a combination of phylogenetic and biochemical analyses, AlphaFold2 structure prediction, molecular dynamics simulations, and mutagenesis. We discovered an unusually narrow transient access path to the central cavity of MtABCG46 that constitutes an initial filter responsible for the selective translocation of these phenylpropanoids through a lipid bilayer. Furthermore, we identified remote residue F562 as pivotal for maintaining the stability of this filter. The determination of individual amino acids that’ impact the selective transport of specialized metabolites may provide new opportunities associated with ABCGs being of interest, as a clinically relevant group of proteins.

## Introduction

ATP-binding cassette (ABC) transporters are prominent proteins that translocate molecules through biological membranes using ATP as a source of energy^1^. Members of the ABC family are common in all domains of life, but plant genomes are exceptionally rich in genes encoding them^2,3^. Taking into account their structure and phylogenetic relationships, most ABC proteins have been classified into eight subfamilies, designated ABCA-ABCH^4^. The particularly numerous plant ABC transporters translocate diverse molecules, such as lipids, phytohormones, carboxylates, heavy metals, chlorophyll catabolites, and xenobiotic conjugates, across various biological membranes^5^. In this manner, they participate in diverse processes, including organ growth, nutrition, development, responses to abiotic stresses, as well as both symbiotic and antagonistic relationships^2,6,7^.

Especially abundant in plants are genes encoding so-called full-size ABCG transporters. A full-size ABCG transporter is a single polypeptide forming two transmembrane domains (TMDs), which constitute a membrane-spanning region, and two cytosolic domains called nucleotide-binding domains (NBDs). Depending on the transporter subfamilies, single TMD has 5 to 10 transmembrane α-helices. Full-size members of the G family are characteristic for plants and fungi and distinct in their reverse organization of domains in the subunits namely NBD1-TMD1-NBD2-TMD2^4,8^. Moreover, in comparison to thoroughly studied ABCB proteins, where α-helices forming TMDs represent a so-called domain swap arrangement, ABCG transporters revealed a different TMD organization, in which not individual helices but the entire TMD rotates as a solid body during the transport of molecules^9^. Full-size ABCGs were initially identified in *Saccharomyces cerevisiae* and the clinically relevant fungus, *Candida albicans*. They were described as Pleiotropic Drug Resistance (PDR) proteins because they can act as efflux pumps, removing diverse molecules from these unicellular organisms, including exogenously applied drugs used in medical treatments, thus conferring resistance against large sets of chemicals^10^. Consequently, PDR proteins have attracted interest of biotechnologists and medical scientists. However, despite strenuous efforts we still have limited understanding of their *modus operandi*.

Characterized plant full-size ABCGs include proteins that can transport several molecules that are not necessarily related but are usually endogenous metabolites. For instance, ABCG37 of *Arabidopsis thaliana* is involved in translocation of the auxin precursor, indolyl-3-butyric acid (IBA)^11^, and the phenolic compound scopoletin^12,13^. Arabidopsis ABCG36/PEN3/PDR8 transports, *inter alia*, IBA, the Brassicales-specific phytoalexin camalexin, heavy metals, and possibly monolignols14–21. However, despite initially considered as functional homologs of yeast multidrug pumps, at least certain plant full-size ABCG proteins appear to be selective towards translocated molecules. The specialization is proposed to be a consequence of a sophisticated chemodiversity exemplified by specialized metabolism that requires tightly controlled distribution of metabolites via dedicated transporters^3,5^.

Phenylpropanoids are a large class of specialized plant metabolites with many important roles in plant biology, medical applications and industrial uses^22^. ABCG46 (formerly known as ABCG10) of the legume *Medicago truncatula*, which is required for efficient *de novo* production of the phenylpropanoid-derived phytoalexin medicarpin, selectively translocates 4-coumarate and liquiritigenin. Notably, structurally similar phenylpropanoids like naringenin, isoliquiritigenin, and 7,4’-dihydroxyflavone are not transported by MtABCG46^23,24^.

Despite the recent progress in structural research regarding ABCG proteins^25–29^, progressing from static atomic structures to an understanding of molecular mechanisms of behind the substrate recognition and transport has been challenging^30^. This is partly because of the difficulty in obtaining experimental data at atomic detail, but also because of the lack of efficient sampling of the intricate process of a complete transport cycle. Fortunately, AlphaFold2^31^ has proven utility for addressing the first of these problems by providing impressively accurate 3D structures of proteins from the amino acid sequence, even for large trans-membrane hydrophobic proteins that are difficult to crystalize^32^. Structurally, although the complete transport process is still elusive, it is accepted that ABCG exporters start this process with an inward-facing (IF) conformation that allows substrate migration from the cytosolic environment to the central cavity. The protein then undergoes large structural rearrangements to an outward-facing (OF) conformation, enabling release of the substrate to the extracellular environment. Binding, hydrolysis and release of ATP as well as substrate recognition followed by its migration contribute to the intricacy of the overall transport process^25,29^.

Using a combination of phylogenetic and biochemical analyses, AlphaFold2 structure prediction, molecular dynamics simulations, and mutagenesis we have identified a transient access path in MtABCG46 that is directly involved in the recognition and passage of 4-coumarate and liquiritigenin through the plasma membrane. Moreover, we have identified F562 as a critical residue for the architecture of this access path responsible for selective transport.

## Materials and methods

### Plant material

*Nicotiana tabacum* Bright Yellow 2 (BY2) suspension cell cultures^53^ were grown in Murashige and Skoog medium supplemented with 2.72 mM KH_2_PO_4_, 0.56 mM myoinositol, 3 μM thiamine, 0.9 μM 2,4-dichlorophenoxyacetic acid, and 87.64 mM sucrose, in the dark at 26 °C on an orbital shaker (130 rpm), and diluted 1:5 every week.

### Genetic constructs

All genetic constructs were based on the pMDC43 vector, carrying a GFP tag sequence^54^. Particular mutants were prepared from a p35S::GFP-MtABCG46 construct by GenScript. All plasmid constructs were confirmed by DNA sequencing.

### Plant transformation

Stably transformed BY2 cells were generated by co-cultivation of 5-day-old BY2 suspension cells with *Agrobacterium tumefaciens* strain AGL1^55^ carrying the pMDC43 vector containing a particular variant of the MtABCG46 sequence or the empty pMDC43 vector, as previously described^24^.

### Confocal microscopy

5-day-old suspension cell cultures overexpressing GFP-fused MtABCG46 variants were observed by laser scanning confocal microscopy (with a Leica TCS SP5 AX v.2.7 instrument).

Plasma membranes of sampled cells were stained with FM4-46 (ThermoFisher Scientific). Obtained pictures were analyzed using Leica LAS AF software. Fluorescent signals from GFP and FM4-64 were pseudo-coloured in green and magenta, respectively.

### Preparation of plasma membrane vesicles

Microsomal fractions were isolated from 12 g portions of BY2 suspension cell cultures as previously described^56^. Plasma membrane fractions of the microsomal isolates were enriched by partitioning in an aqueous two-phase system, also as previously described^57^. The quality of obtained microsomes was tested with 9-amino-6-chloro-2-methoxyacridine (ACMA; Invitrogen A1324) fluorescence quenching assays.

### Transport analysis with plasma membrane vesicles

The transport of phenolic compounds uptake was studied by the rapid filtration technique with 4-coumarate, liquiritigenin, isoliquiritigenin, 7,4’-dihydroxyflavone and the plasma membrane microsomes using nitrocellulose filters (0.45 mm pore-size; Millipore). The transport assays were performed with microsomes corresponding to 520 ng·μl^-1^ protein concentration mixed with transport buffer (10 mM Tris-HCl, 10 mM EDTA, 10% sucrose, pH 5.0), the selected phenolic (750 μM), 100 μg·mL^-1^ creatine kinase, 10 mM creatine-phosphate and 1 mM of MgCl2, in the presence and absence of 4 mM ATP. After 3 minutes incubation at 24 °C, 0.3 mL of each reaction mixture was immediately loaded on a prewetted filter and rapidly washed with 10 mL of ice-cold transport buffer. The filters were air-dried for an hour then incubated in 80% MetOH with 0.1% formic acid. Phenolic compounds were extracted by adding chloroform:water mixture (1:0.25 sample volume) to the sample and centrifugation for 30 minutes in 13200 rpm. The dried samples were dissolved in 80% methanol and subjected to HPLC/MS analyses, as described below. For competition assays, tested molecules were added, each at 750 μM, together to the transport buffer. Experiments were repeated three times with independent vesicle preparations unless stated otherwise.

### HPLC/MS analysis

Samples were analyzed by liquid chromatography–electrospray ionization–tandem mass spectrometry (LC/ESI/MS) using a Waters UPLC Acquity system, equipped with a C18 RP column, connected to a Bruker micrOTOF-Q II mass spectrometer. The mobile phase consisted of a gradient of 0.5% formic acid (v/v) in water (A) and 0.5% formic acid (v/v) in acetonitrile (B). The m/z range of the recorded MS spectra was 50–1000. The MS was operated in positive and negative ion modes for phenolics and carboxylic acids, respectively.

### Multiple sequence alignment and data filtering

To select full-size ABCG from multiple groups of plants, we subjected 1KP transcript data to tBLASTn searches using MtABCG46 as a query sequence with an E-value cutoff of 1e-5. Next, 26 889 1KP samples longer than 1000 bp were translated into six frames using software written in Visual C#. The longest ORFs starting with the ATG codon, after excluding the duplicates, were selected and further verified, by a BLASTp search against known ABC transporters belonging to different subfamilies was conducted. 1839 1KP samples were assigned to a full-size ABCG subfamily with over 70% coverage and an E-value of 0.0, and used for the subsequent analysis. Multiple sequence alignment (MSA) of the full-size ABCG amino acid sequences was performed using the MUSCLE algorithm^58^ in MEGA X^59^. For conservation analysis, complete alignment of predicted amino acid sequences of 1576 plant full-size ABCG transporters was submitted to the ConSurf server with default settings^39,40^. We also submitted extracted taxa-specific alignments (66, 205, 91, 388 and 725 green alga, bryophyte, pteridophyte, monocots and core eudicot sequences, respectively) to the ConSurf server, and subjected the complete alignment (1576 sequences) to co-evolution analysis using the Gremlin server^41,42^ with default settings.

### Modeling of the 3D structure

The amino acid sequence of the ABCG46 transporter of *Medicago truncatula* was obtained from the UniProt database (accession no. A0A396JDZ5), and submitted to a local installation of AlphaFold2 v2.1.0^31^. The resulting models were evaluated using the per-residue lDDT-Cα metric^33^ and results from PROCHECK software^60^.

To define positions of the Mg^2+^ ions, experimental structures of ABCG2 from *Homo sapiens* (PDB ID: 6hbu) and Pdr5 from *Saccharomyces cerevisiae* (PDB ID: 7p06) were employed as templates. The NBD region of the model and experimental structures were superimposed using TM-align software^34^, and positions of the Mg^2+^ ions were copied to the model. For the ATP molecules, the NBD regions were also superimposed, but coordinates of the ATP atoms were not used directly. Instead, a docking box enclosing the molecules was built, then AutoDock Vina v1.1.2^61^ was used to determine their most suitable positions. Mutations at residue F562 to alanine, leucine and tyrosine were performed with the tleap module of the Amber20 package^62^, and one system of the wild-type protein was built without ATP and Mg^2+^ ions as an APO variant.

### Molecular Dynamics (MD) simulations

The protein model was protonated with the H++ server^63^ at pH 7.0. The protonated protein was embedded in a 1-palmitoyl-2-oleoyl-sn-glycero-3-phosphocholine (POPC) lipid bilayer using the CHARMM-GUI server^64–68^. For the system, the size of the simulation box was set to 130 Å, with water thickness of 15 Å, and KCl salt concentration of 0.1 M. The water model employed was OPC^69^, and the HMR^70^ method was used to enable 4 fs simulation timesteps. The MD engine employed was Amber20^62^ with the pmemd GPU implementation^71^, the ff19SB force field^72^ was used for the protein, lipid17 for the POPC membrane, and previously presented parameters for the ATP molecules^73^. The system was minimized with 2500 steps of steepest descent, followed by 2500 steps of conjugated gradient, applying only positional restraints to the protein, ATP, and Mg^2+^ ions, but positional and dihedral restraints to the membrane (Table S3). After the minimization, a series of short equilibration steps was applied to increase temperature and release restraints in the system, each consisting of 1 ns of simulation time. After the equilibration stage, an extra 100 ns equilibration step was applied in which restraints were only maintained for the Mg^2+^ and ATP molecules. Finally, 100 ns unrestrained MD to fully equilibrate the system was followed by a production phase of 400 ns unrestrained MD. For every variant considered, five replicas of the production phase were simulated.

### Access path detection, classification, and selection

For path computation and detection, CAVER v3.0 software^35^ was employed with a 0.9 Å probe radius, 6 Å shell radius, and 4 Å shell depth. For clustering, the average-link hierarchical algorithm was used, the maximum number of clusters was set to 50, and clustering threshold to 3.5. The starting position of the CAVER calculation was set to employ the center of mass of residues 562, 566, 1213 and 1217. Subsequently, TransportTools software v0.9.0^36^ was used to obtain a comprehensive and comparative view of the path network across all variants. In TransportTools, the clustering method was set to complete with a clustering cutoff of 0.5 Å for the analysis of the wild-type protein, then switched to the average method with a clustering cutoff of 2.0 Å for the comparison of all variants simultaneously, with all other parameters left as default. The candidate selected as the correct access path was further sorted by the bottleneck radius and the 100 widest paths of each variant were used in the analysis and further ligand migration experiments.

### Ligand migration and energy barrier calculation

To study the capability of each variant to transport the four tested phenolic compounds, molecular docking across selected paths was performed with CaverDock software v1.1^46^. 3D structures of the ligands of the four compounds were obtained from the PubChem database: liquiritigenin (CID 114829), isoliquiritigenin (CID 638278), 4-coumarate (CID 637542) and 7,4’-dihydroxyflavone (CID 5282073), then the MGLTools v1.5.6^74^ was used to prepare the files for CaverDock with the prepare_ligand4.py script with the default settings. In the same way, all the snapshots of the MDs from the variants where a path was selected were processed with the prepare_receptor4.py script with default settings. The exhaustiveness for CaverDock runs was set to one, using a single CPU core per task and eight parallel workers per combination of variant-path-ligand. Finally, the results were analyzed with in-house Python scripts.

### Auxiliary analyses of data from MD simulations

Using the cpptraj^75^ module of the Amber20 package, initially the heavy atoms’ root mean square deviation (RMSD) and the root mean square fluctuation (RMSF) of the whole system were calculated. However, since the main focus was the TMD region, separate RMSDs for specific regions were also calculated (Fig. S3 and Table S4). For the membrane, the lipid order parameters of lipid tails were evaluated together with the mass density with cpptraj to ensure the proper equilibrium behavior of the POPC membrane. To evaluate the behavior of the TMD helices forming the candidate access path, α helices 2, 5, 8, and 11 were analyzed with the HELANAL module of MDAnalysis v2.0.0^76,77^.

### Statistical analyses

Statistical analyses were performed using GraphPad Prism software v9.0. The normality of distribution assumption was assessed for particular groups of values by Anderson-Darling, Shapiro-Wilk, and Kolmogorov-Smirnov normality tests. If data did not meet normal distribution criteria, non-parametric tests were applied (the two-tailed Mann-Whitney test or Kruskal-Wallis test with post hoc Dunn’s multiple comparison test). *P* values obtained can be found in the Supporting Statistical Data.

## Results

### The MtABCG46 model features an unusually occluded central cavity connected with the intracellular environment by a transient access path

The MtABCG46 model obtained from AlphaFold2 has good overall quality according to the per-residue lDDT-Cα metric^33^, with limited confidence for only a few regions (Fig. S1). Moreover, these regions correspond to highly dynamic motifs in related ABCG proteins^25,27,29^, for which single structures are hard to define, attesting to high model quality. According to the template modeling score (TM-score)^34^, the model has a very similar fold to the recent full-size PDR5 structure of *Saccharomyces cerevisiae* (ScPDR5)^29^, human ABCG1 (HsABCG1)^27^, and half-size human ABCG2 (HsABCG2)^25^ with TM-scores of 0.82, 0.72, and 0.66, respectively (Fig. S2). In the structure, the typical arrangement of domains in a full-size ABCG transporter is clearly visible (Fig. S3), with TMD regions, each composed of six α-helices, and the two half-size domains joined by a ~55-residue linker region. A comparison of central cavities of MtABCG46 and ScPDR5 revealed a small disconnected cavity in the TMD region of MtABCG46 (Fig. 1a). In contrast, a wide-open cavity in the TMD region that directly connects with the cytosolic environment was observed in ScPDR5 (Fig. 1b).

**Fig. 1.**
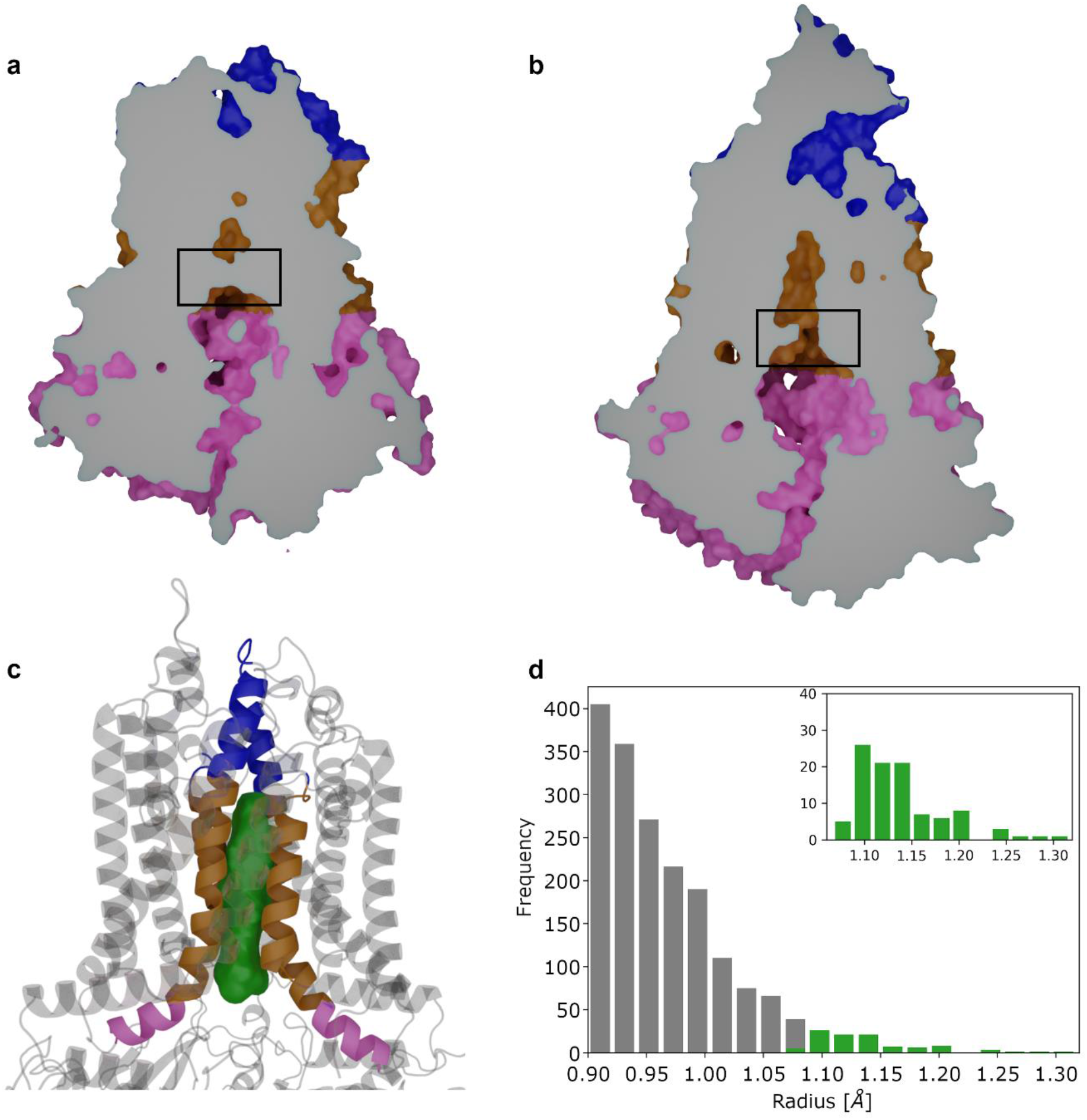
Structure and dynamics of the central cavity and its access path in MtABCG46. **a** Sliced surface side view of the MtABCG46 model obtained from AlphaFold2 featuring an occluded central cavity (the closure indicated by a black rectangle). **b** Sliced surface side view of a cryo-EM structure of ScPDR5 (PDB ID: 7p04) with an open cavity (indicated by a black rectangle). **c** Structural representation of part of MtABCG46, showing the overall volume of the access path ensemble (green) and TMD helices forming it (intra-cellular region in pink, trans-membrane region in gold, and extra-cellular region in blue). **d** Bottleneck radius distribution of access path ensemble connecting the cavity in MD simulations of MtABCG46, the widest 100 paths are colored green and shown in the inset.

To explore possible paths enabling access of cognate ligands into the central cavity, we analyzed molecular dynamics (MD) simulations for temporarily opened continuous internal voids formed within the MtABCG46 structure using CAVER 3.0^35^ and TransportTools^36^. This revealed a network of putative transport paths from the bottom of the cavity to the intracellular, extracellular and membrane regions (Fig. S4a). To help identification of the likeliest localization of a functionally relevant entrance to such a path along the membrane’s Z-axis, we ran six independent MD simulations of systems composed of the endogenous MtABCG46 substrate, liquiritigenin, and a membrane. These simulations showed that liquiritigenin preferentially stayed between the heads and tails of the membrane phospholipids (Fig. S4b, c). Consequently, among all paths that opened in this region, the most prevalent (open for ~ 9 % of the total simulation time) was the third-ranked one (Table S1) with an average bottleneck radius of 0.97+0.06 Å (Fig. 1d). This path provided the most straightforward access to the central cavity from the intracellular region (Fig. 1d and Fig. S4d). Interestingly, an equivalent path was practically undetectable in simulations of MtABCG46 without bound ATP molecules (Fig. S5). The remaining paths leading to the intracellular region were not considered due to their much lower frequency and much longer, curved geometry (Table S1).

### Phylogenetic analyses of residues of the MtABCG46 central cavity

To investigate the importance of the architecture of internal voids in MtABCG46 for the passage of specific phenylpropanoids, we mapped residues contributing to the surface of the access path and central cavity during the MD simulations (Fig. S6). We hypothesized that variations in the recognition and transport of diverse molecules by various ABCG proteins may arise from differences in the amino acid sequences that form the cavity. Based on this assumption we generated multiple sequence alignment (MSA) of predicted amino acid sequences of 1839 plant full-size ABCG transporters extracted from the 1 KP project^37,38^, and analyzed it using the ConSurf^39,40^ and Gremlin^41,42^ servers. We then selected residues that contribute to the central cavity, are not fully conserved (ConSurf grade ≤ 8), and display variability that could not be readily explained by co-evolutional links with other residues (Fig. 2a).

**Fig. 2.**
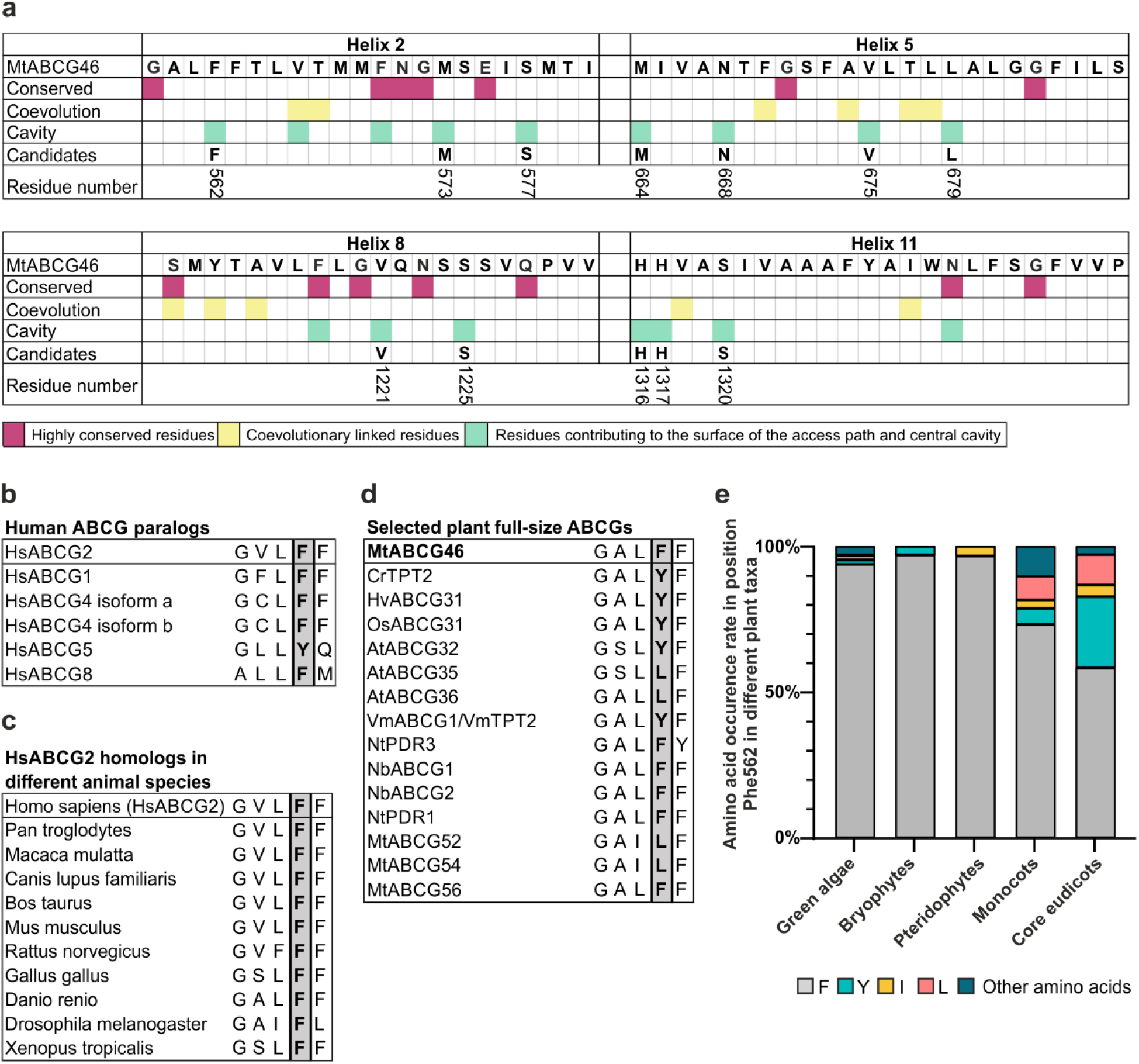
Residue selection for the site-directed MtABCG46 mutagenesis. **a** Schematic representation of candidate residues in sequences of MtABCG46 transmembrane helices 2, 5, 8 and 11 selected using conservation degrees (with Consurf), coevolutionary parameters (with Gremlin) and the structural data. Candidate residues numbered in accordance with the MtABCG46 sequence. **b, c** Alignment of the region corresponding to surroundings of F431 in HsABCG2 for human ABCGs (**b**) and animal homologs (**c**). **d** Alignment of the region corresponding to surroundings of F562 in MtABCG46 for selected plant full-size ABCG transporters. **e** Frequencies of occurrence of indicated amino acids in the residue corresponding to MtABCG46 F562 in full-size ABCG sequences of indicated taxa.

One of the selected amino acids, residue F562 in TMD helix 2, particularly drew our attention due to its correspondence to F431 in HsABCG2 (Fig. 2 and Fig. S7), a putatively important residue for ligand recognition and binding^43^. Residue F431 is highly conserved in human ABCG transporters and fully conserved among ABCG2 homologs in several animal species (Fig. 2b, c). However, in plant full-size ABCG transporters, variations at the F562 position include amino acids such as tyrosine, leucine, and isoleucine (Fig. 2d). Phylogenetic analyses revealed that the prevalence of amino acids other than phenylalanine at this position is significantly higher in monocots and eudicots than in non-seed plants (Fig. 2e). Notably, such expansion of variability in angiosperms was not observed for other residues that directly contribute to the cavity surface (Fig. S8). Since angiosperms have a higher degree of chemodiversity (and associated specialized metabolism) this led to the conclusion that variability at this position might be meaningful for plant ABCG transporters and/or possibly MtABCG46 selectivity.

### F562 substitutions profoundly affect the selectivity of MtABCG46-mediated transport

To address the importance of F562 for MtABCG46-mediated transport of phenylpropanoids, we substituted it for the other two most frequent amino acids at this position in plant full-size ABCG transporters, tyrosine and leucine (Fig. 2e), as well as alanine. Alanine is often used in such analyses as it eliminates the side-chain beyond the β carbon, but does not alter the main-chain conformation^44^.

All our MtABCG46 variants, including the native form (hereafter: wild type, WT), tagged with GFP at the N terminus, were introduced into BY2 tobacco suspension cell cultures, a well-established heterologous expression systems for biochemical studies of ABCG proteins^8,45^. To confirm the presence and correct localization in the plasma membrane (PM) of the transporters in BY2 lines, the colocalization of GFP-tagged MtABCG46 variants with a PM marker, FM4-64, was checked with confocal microscopy (Fig. 3a).

**Fig. 3.**
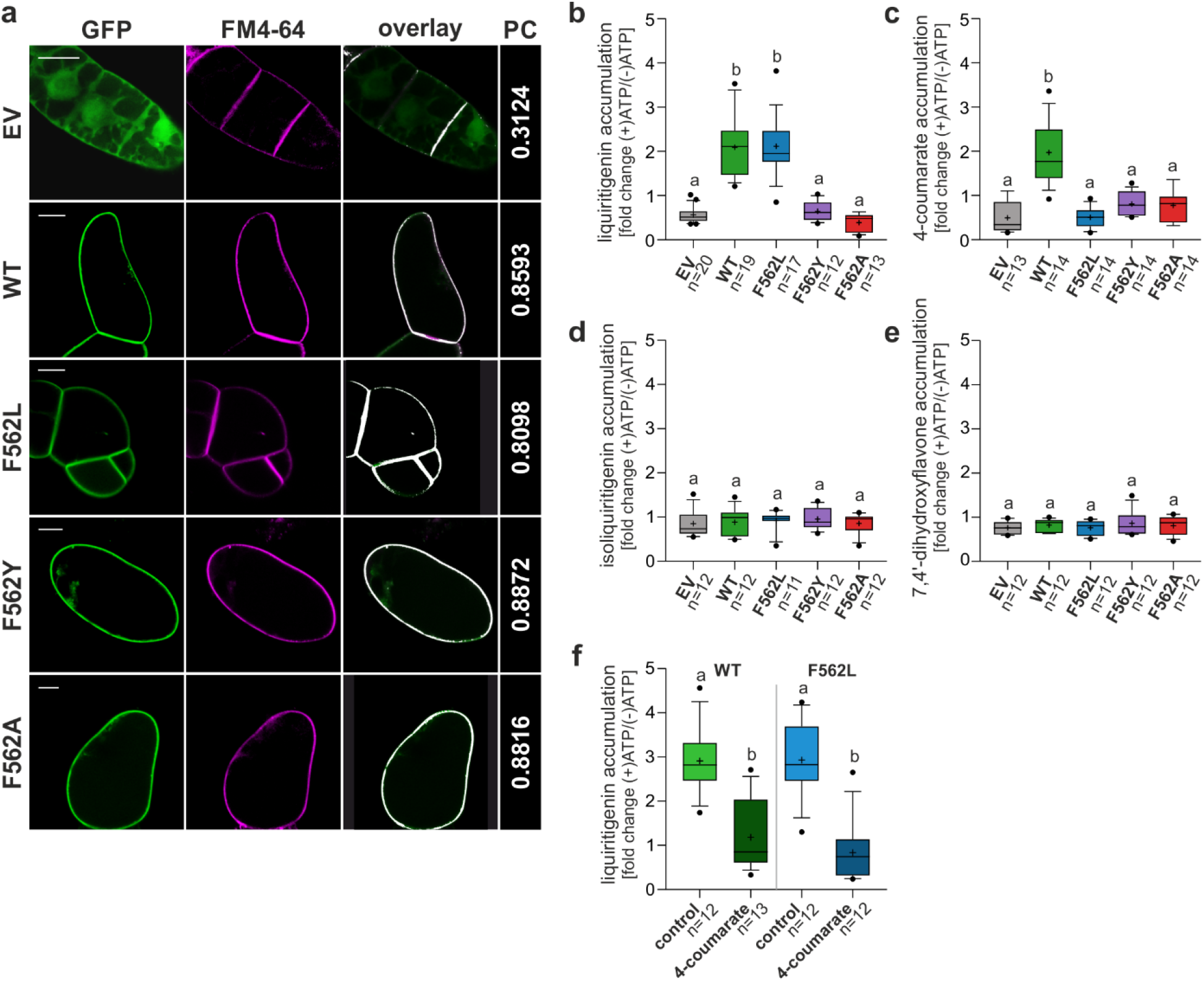
Transport assays with microsomes derived from BY2 suspension cell cultures. **a** Plasma membrane localization of the MtABCG46 variants tagged with GFP: WT, F562L, F562A, and F562Y in BY2 suspension cell cultures. Images with GFP and FM4-64 fluorescence pseudo-coloured in green and magenta, respectively. PC - Pearson’s correlation coefficients for the colocalization of GFP and FM4-64 in the plasma membrane, visualized as an overlay. Scale bars, 20 μm. **b-e** Transport of liquiritigenin (**b**), 4-coumarate (**c**), isoliquiritigenin (**d**) and 7,4’-dihydroxyflavone (**e**) in microsomes derived from BY2 suspension cell cultures expressing empty vector (gray) or indicated variants of MtABCG46: WT (green), F562L (blue), F562Y (purple) and F562A (red). **f** Competitive transport of liquiritigenin and 4-coumarate in microsomes expressing WT and F562L MtABCG46: means ±SD from three biological replications, each with at least three technical replications. **b-f** Values are presented as fold change between (+)ATP and (-)ATP as a control. In each box-and-whiskers plot: the central black line indicates the median; ‘+’ indicates the mean; the box extends from the 25th to 75th percentile; the whiskers extend from the 10th to the 90th percentile, and points below and above the whiskers are marked by individual dots. Different lowercase letters indicate significant differences: *P* < 0.001 (**b, f**); *P* < 0.01 (**c**), *P* < 0.05 (**d**, **e**). P values, determined by the Kruskal–Wallis test with a post hoc Dunn’s multiple comparison test (**b-e**) or Mann-Whitney test (**f**) can be found in Supporting Statistical Data.

Effects of mutations were investigated by ATP-dependent, MtABCG46-mediated, transport assays of liquiritigenin, 4-coumarate, isoliquiritigenin, and 7,4’-dihydroxyflavone into PM inside-out vesicles derived from BY2 lines. Consistent with previous observations^24^, liquiritigenin and 4-coumarate accumulated in vesicles from lines expressing WT MtABCG46 (Fig. 3b, c) but not isoliquiritigenin or 7,4’-dihydroxyflavone (Fig. 3d, e). Further experiments with lines expressing variants of MtABCG46 revealed that F562Y and F562A substitutions abolished transport of liquiritigenin and 4-coumarate (Fig. 3b, c). None of the variants were able to translocate isoliquiritigenin nor 7,4’dihydroxyflavone (Fig. 3d, e). Interestingly, the F562L variant, accumulated liquiritigenin similarly to the WT MtABCG46, however, it was not able to transport 4-coumarate (Fig. 3b, c).

Previous research has shown that liquiritigenin is a competitor of 4-coumarate in MtABCG46-mediated transport^24^. Analyses of liquiritigenin transport in the presence of 4-coumarate confirmed that this effect is mutual, i.e., 4-coumarate also reduces the rate of MtABCG46-dependent liquiritigenin transport (Fig. 3f). Experiments also revealed that although F562L cannot transport 4-coumarate, the latter is still a competitor for liquiritigenin (Fig. 3f).

### F562 substitutions affect the viability of the transient access path through rearranged TMD helices

In MD simulations of our four variants (WT, F562L, F562Y, and F562A) there were no significant differences in stability or flexibility between the mutant and WT proteins (Figs. S9-12). However, there were considerable differences in the availability of open access paths leading to the central cavity, which were lower in all mutants than in the WT (Table S2 and Fig. S13a). The 100 widest paths were almost as wide in the F562L mutant as in WT, but they were markedly narrower in F562Y and F562A mutants (Fig. S13b), mainly because they had more constricted entrances (Fig. 4a, Fig. S14), presumably hampering access of bulky molecules to the central cavity. Accordingly, we hypothesized that mutation of residue F562 affects the structural arrangement of residues forming the access path, thereby significantly reducing its overall transport capability.

**Fig. 4.**
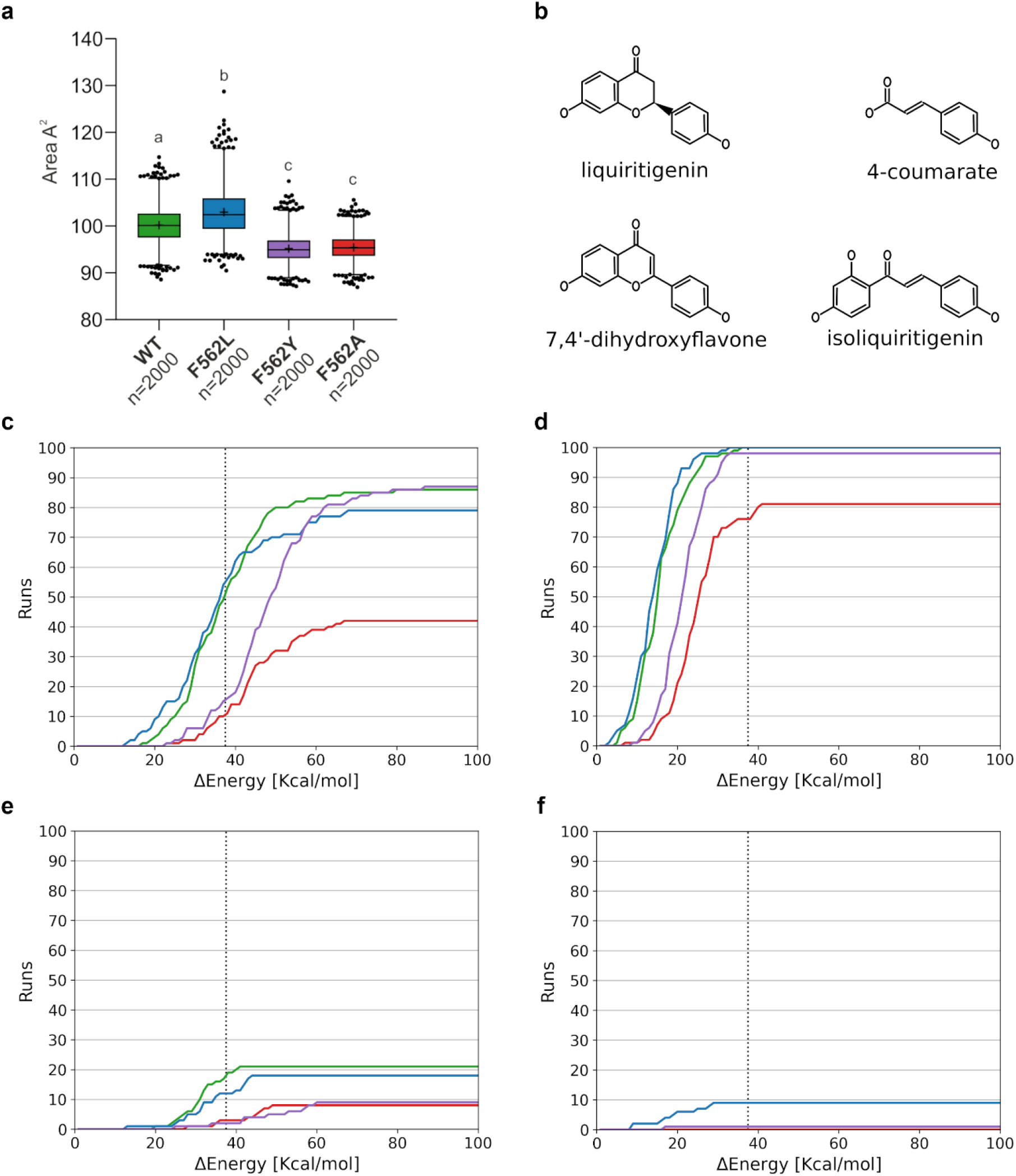
Effects of F562 mutations on the accessibility of the path to the central cavity of MtABCG46. **a** Area of the entrance to the access path in MtABCG46. For the definition of the area, see Fig. S14. In each box-and-whiskers plot: the center black line indicates the median; ‘+’ indicates the mean; the box extends from the 25th to 75th percentile; the whiskers extend from the 1st to the 99^th^ percentile, points below and above the whiskers are marked as individual dots. Different lowercase letters indicate significant differences, *P* < 0.0001. P values, determined by the Kruskal–Wallis test with a post hoc Dunn’s multiple comparison test can be found in Supporting Statistical Data **b** Structures of ligands evaluated with CaverDock. **c-f** Cumulative distributions of energetic barriers for transport to the internal cavity for the widest 100 paths for liquiritigenin **(c)**, 4-coumarate **(d)**, 7,4’-dihydroxyflavone (**e**), and isoliquiritigenin **(f)**. In **c-f** curves obtained from experiments with WT, F562L, F562Y, and F562A variants are marked in green, blue, purple, and red, respectively, and the median value for WT-liquiritigenin transport is indicated by a black dashed vertical line.

To address this possibility, we assessed the viability of the 100 widest paths in each variant for the access of liquiritigenin and 4-coumarate, as well as isoliquiritigenin and 7,4’-dihydroxyflavone, which are chemically similar but not transported by MtABCG46 (Fig. 4b), from the intracellular environment into the central cavity using CaverDock^46^. Each run of CaverDock yields an energetic profile of the binding energy between a ligand and protein along the path, which can be translated as an energetic barrier that must be overcome for a ligand to reach the central cavity (Fig. S15). Such an approach has proven utility in detecting hotspot residues for protein engineering^47^, and correlating energetic barriers with biochemical rates^48,49^. The cumulative distribution of the energy barriers for all variants showed that liquiritigenin could reach the central cavity equally well in WT and F562L mutant (with a median energy barrier of 38.5 kcal·mol^-1^) (Fig. 4c). Frequencies of successful migration events in F562Y and F562A were much lower: 16% and 11% at the median WT energy barrier, respectively (Fig. 4c). Moreover, to reach the same frequency of successful access to the central cavity, liquiritigenin had to overcome energy barriers of up to 50 kcal·mol^−1^ in F562Y and this frequency could not be reached at all in F562A (Fig. 4c). In contrast, 4-coumarate reached the central cavity in almost all simulated migration events within this threshold in all variants (Fig. 4d). Such a surprising discrepancy with overall transport assays of this substrate indicates that mutations at F562 also affect transport of the substrates after they have reached the central cavity. Conversely, isoliquiritigenin and 7,4’dihydroxyflavone rarely reached the cavity in any variant (Fig. 4e, f).

Structurally, the introduced modifications of residue 562 disrupted non-covalent contacts (Fig. S16), changing the bending and twisting angles of the TMD helices forming the access path (Figs. S17, S18). In WT, residue F562 maintained a parallel displaced π-stacking interaction with residue F684 during the simulations (Fig. S16a), a feature shared with the F562Y mutant (Fig. S16c). Loss of this non-covalent contact in F562A resulted in a considerable displacement of TMD helix 5 (Fig. S16d). F562L also lost the π-stacking interaction, but maintained stability of this helix (Fig. S16b), likely due to remaining Van der Waals interactions of its sidechain. Interestingly, the displacement of TMD helix 5 in F562A allowed marked bending and to some extent twisting of TMD helix 8 (Fig. S17c, S18c). Furthermore, the bending of TMD helices affected not only the structure of the central cavity but also the path entrance, narrowing the available space for molecules to enter (Fig. 4a). This reduction of available space at the entrance to the access path was observed in F562A and F562Y mutants, although no TMD helix deformations were observed in the F562Y mutant. The additionally introduced hydroxyl group in F562Y formed an H-bond with Y1213 (in TMD helix 8) during the whole simulation time. This pulled the modified residue and Y1213 closer to each other, thereby rearranging the orientation of the neighboring residues in TMD helix 8 and changing its twist helical angle (Fig. S18c). Moreover, residue Y1213 formed an H-bond with N1331, a highly conserved residue (Fig. 2a), in all performed MD simulations. Hence, disturbance of the normal behavior of these residues could influence ligands’ binding in the central cavity.

## Discussion

AlphaFold2 has recently proposed suitability for modelling ABCG proteins, with the ability to provide similar levels of accuracy as for soluble proteins^32^. The MtABCG46 structure we obtained using AlphaFold2 has the typical architecture of full-size ABCG transporters described in the literature^50^. Moreover, our MD simulations identified transiently formed access paths to the central cavity from the intracellular region (Fig. 1 and Table S1). This cavity and its access paths were localized in a similar region to the open cavities observed in HsABCG1^27^, HsABCG2^25^, and a recently obtained cryo-EM structure, ScPDR5^29^ (Fig. S19). Several substrates (such as rhodamine 6G, cholesterol, and mitoxantrone) have been found to occupy equivalent regions of ScPDR5, HsABCG1, and HsABCG2, respectively^25,27,29^, indicating the potential importance of this access path and central cavity of MtABCG46 for the transport.

Tested substitutions of the selected residue F562 in the central cavity (F562L, F562Y, and F562A) severely affected MtABCG46-mediated transport of 4-coumarate and liquiritigenin, highlighting this residue’s importance for phenylpropanoid transport in *M. truncatula*. Transport assays of four phenylpropanoids with the four MtABCG46 variants showed that none of them can transport isoliquiritiginen or 7,4’dihydroxyflavone (Fig. 3d, e), in accordance with indications from CaverDock analyses that likelihoods of these compounds accessing the central cavity are extremely low (Fig. 4e, f). We found no significant difference in the energy barrier for liquiritigenin entry into the central cavities of WT and F562L MtABCG46, which is sufficiently low for successful transport of this substrate by both variants. In contrast, liquiritigenin could not enter the cavity of F562A or F562Y, probably due to a collapse of the access path caused by their mutations (Fig. 4a), which consequently lack ability to transport it (Fig. 4c, 3b). Thus, our data indicate that restriction of access to the central cavity is a major contributor to substrate selectivity in early stages of the transport process.

Curiously, we found that only MtABCG46 WT could transport 4-coumarate (Fig. 3c). CaverDock analysis detected no difference in accessibility of the central cavities of MtABCG46 WT and F562L for this metabolite, and only limited disadvantage in the other two mutants (Fig. 4d). Nevertheless, 4-coumarate showed similar competitive behavior towards liquiritigenin in F562L to that observed in WT (Fig. 3f). This suggests that 4-coumarate may still be able to reach the central cavity, and the transport failure may be linked to perturbation in subsequent stages of the transport cycle, *e.g*., recognition of the substrate bound in the central cavity or its stabilization during the subsequent conformational change (from IF to OF) of the transporter. To explore this possibility, we investigated the availability of polar interactions provided by residues of the central cavity to 4-coumarate. We found only three polar residues around the deepest part of the central cavity: Y1213, T1214 and N1331. Of these, T1214 has limited accessibility as it is oriented outwards of the cavity, and Y1213 maintains constant H-bond contact with the ketone group of N1331 in all MD simulations. Hence, the amine group of N1331 is the only one to act as a hydrogen donor. Notably, a high conservation score was obtained for residue N1331 in our MSA analysis (ConSurf grade 9), indicating that it has functional importance. In our MD simulations, all investigated mutations of residue F562 resulted in a considerable change in the conformation of N1331 (Fig. S20) perturbing the putative interaction between N1331 and 4-coumarate, and possibly hampering transport of the latter.

In general, the molecular determinants of sequence changes resulting from protein adaptation to various biochemical needs remain obscure. Our experimental data, together with phylogenetic observations, suggest that adaptation of ABCG proteins associated with evolutionary pressures in plants has resulted in a variability of some key residues, such as F562 (Fig. 2e). Such variability is essential for transporters to fulfill their functional roles in diverse, complex chemical scenarios. ABCG protein own a recognized role in secretion of end-products of specialized metabolism, which may have valuable biotechnological applications. Notably, specialized metabolites of plants are potential sources of novel substances that may help efforts to overcome the resistance of clinically relevant pathogenic fungi to commonly applied classes of drugs, such as fluorinated pyrimidine analogs, polyenes, allylamines, and azoles^51^.

On the other hand, despite strenuous efforts to elucidate structures and functions of transporters involved in drug resistance, example of being CDR1 from *C. albicans*, including extensive mutational and suppressor analyses, we have poor understanding of their *modus operandi*^52^. As new drugs are often specialized metabolites of plants, or their derivatives, analysis including exploration of the relations of their translocation patterns with endogenous ABCG transporters is a promising alternative approach. We foresee sequence-structure-dynamics exploration fueled by AplhaFold2 presented here as an alternative mean to overcome limitations in structural studies of membrane transporters, which can help to identify residues that define functional properties of that important subfamily of ABC transporters.

## Supporting information

Supplemetary Statistical Data

## Supporting Information

Supporting Figures and Tables (PDF) include details on the MtABCG46 model; the localization of access pathways leading into central cavity and description of their characteristics; structural and sequential alignment of MtABCG46 and HsABCG2; conservation score of the inner cavity forming residues; RMSD and RMSF of MtABCG46 variants observed in simulations; structural coupling between access path and transmembrane helices; interactions between key residues of transmembrane helices; the description of perturbation of transmembrane helices by mutagenesis; comparison of internal cavities among related ABC transporters; the description of conformational space adopted by N1331 in MtABCG46 variants observed in simulations; duration and restraints applied in individual simulation stages. Additionally, the results of statistical analyses are summarized in the Supporting Statistical Data (XLSX).

## Statements and Declarations

### Author Contributions

M.J. devised and supervised the project. J.Br. designed, interpreted and supervised the computational experiments. K.P. and M.G. selected residues for the site-directed mutagenesis (conservation and coevolution analyses). C.S.B. performed, analyzed, and interpreted results of computational experiments. W.B.L. and K.P. designed, performed and interpreted results of the transport experiments (BY2 transformation, isolation of microsomes, transport and competition assays). A.P. contributed to BY2 transformation, isolation of microsomes, transport and competition assays. W.B.L was responsible for HPLC/MS analysis and statistical analysis. J.Ba., A.B., M.R. ABCG selected sequences acquired in the 1KP project. J.Ba. and W.B.L., performed microscopic observation. C.S.B., W.B.L. and K.P. prepared figures. K.P., C.S.B, W.B.L., J.Br., and M.J. wrote the manuscript with the help of the co-authors. All authors discussed and approved the manuscript.

### Data availability

Sequence data reported in this article can be found in the NCBI database under the following accession numbers: MtABCG46 (Medtr_2g102670), MtABCG52 (Medtr_8g014360), MtABCG54 (Medtr_5g070320), MtABCG56 (Medtr_2g101090), AtABCG32 (AT2G26910), AtABCG35 (AT1G15210), AtABCG36 (AT1G59870), NbABCG1a (BAR94041), NbABCG2b (BAR94044), NtPDR3 (Q5W274), NtPDR1 (NP_001312599), HvABCG31/EIBI1 (BAK52288), OsABCG31 (Os01g0177900), CrTPT2 (KC511771), VmABCG1/VmTPT2 (KC511773), HsABCG1 isoform X1 XP_011528108, HsABCG2 NP_004818.2, HsABCG4 isoform a NP_001335120.1, HsABCG4 isoform b NP_001335121.1, HsABCG5 NP_071881.1, HsABCG8 NP_071882.1; HsABCG2 homologs: *Homo sapiens* NP_004818.2, *Pan troglodytes* XP_526633.3, *Macaca mulatta* NP_001028091.1, *Canis lupus familiaris* NP_001041486.1, *Bos taurus* NP_001032555.2, *Mus musculus* NP_036050.1, *Rattus norvegicus* NP_852046.1, *Gallus* XP_421638.4, *Danio rerio* NP_001036240.1, *Drosophila melanogaster* NP_001039227.1, *Xenopus tropicalis* NP_476787.1. The sequences from 1KP analysis, predicted AlphaFold2 models, parameters and input files for MD simulations of all MtABCG46 variants, as well as key restart files, results of tunnel geometry and ligand-migration analyses are available at https://doi.org/10.5281/zenodo.7002738.

## Acknowledgements

The computations were performed at the Poznan Supercomputing and Networking Center.

We thank Dr. Mariusz Czarnocki-Cieciura (IIMCB Warsaw) for obtaining structural predictions from the local installation of AlphaFold2, enabling predictions using the full-length sequence of MtABCG46.

## References

1. Rees, D. C., Johnson, E. & Lewinson, O. ABC transporters: The power to change. Nature Reviews Molecular Cell Biology 10, 218–227 (2009).

2. Hwang, J. U. et al. Plant ABC transporters enable many unique aspects of a terrestrial plant’s lifestyle. Molecular Plant 9, 338–355 (2016).

3. Banasiak, J. & Jasiński, M. ATP-binding cassette transporters in nonmodel plants. New Phytologist 233, 1597–1612 (2022).

4. Verrier, P. J. et al. Plant ABC proteins - a unified nomenclature and updated inventory. Trends in Plant Science 13, 151–159 (2008).

5. Lefèvre, F. & Boutry, M. Towards identification of the substrates of ATP-binding cassette transporters. Plant Physiology 178, 18–39 (2018).

6. Do, T. H. T., Martinoia, E., Lee, Y. & Hwang, J.-U. 2021 update on ATP-binding cassette (ABC) transporters: How they meet the needs of plants. Plant Physiology 187, 1876–1892 (2021).

7. Do, T. H. T., Martinoia, E. & Lee, Y. Functions of ABC transporters in plant growth and development. Current Opinion in Plant Biology 41 32–38 (2018).

8. Kang, J. et al. PDR-type ABC transporter mediates cellular uptake of the phytohormone abscisic acid. Proceedings of the National Academy of Science USA 107, 2355–2360 (2010).

9. Ford, R. C. & Beis, K. Learning the ABCs one at a time: Structure and mechanism of ABC transporters. Biochemical Society Transactions 47, 23–36 (2019).

10. Prasad, R. & Goffeau, A. Yeast ATP-binding cassette transporters conferring multidrug resistance. Annual Review of Microbiology 66, 39–63 (2012).

11. Ito, H. & Gray, W. M. A gain-of-function mutation in the Arabidopsis pleiotropic drug resistance transporter PDR9 confers resistance to auxinic herbicides. Plant Physiology 142, 63–74 (2006).

12. Fourcroy, P. et al. Involvement of the ABCG37 transporter in secretion of scopoletin and derivatives by Arabidopsis roots in response to iron deficiency. New Phytologist 201, 155–167 (2014).

13. Ziegler, J., Schmidt, S., Strehmel, N., Scheel, D. & Abel, S. Arabidopsis transporter ABCG37/PDR9 contributes primarily highly oxygenated coumarins to root exudation. Scientific Reports 7, 3704 (2017).

14. Aryal, B. et al. ABCG36/PEN3/PDR8 is an exporter of the auxin precursor, indole-3-butyric acid, and involved in auxin-controlled development. Frontiers in Plant Science 10, (2019).

15. Matern, A. et al. A substrate of the ABC transporter PEN3 stimulates bacterial flagellin (flg22)-induced callose deposition in Arabidopsis thaliana. Journal of Biological Chemistry 294, 6857–6870 (2019).

16. Strader, L. C. & Bartel, B. The Arabidopsis PLEIOTROPIC DRUG RESISTANCE8/ABCG36 ATP binding cassette transporter modulates sensitivity to the auxin precursor indole-3-butyric acid. Plant Cell 21, 1992–2007 (2009).

17. Stein, M. et al. Arabidopsis PEN3/PDR8, an ATP binding cassette transporter, contributes to nonhost resistance to inappropriate pathogens that enter by direct penetration. The Plant Cell 18, 731–746 (2006).

18. Lu, X. et al. Mutant allele-specific uncoupling of penetration3 functions reveals engagement of the ATP-binding cassette transporter in distinct tryptophan metabolic pathways. Plant Physiology 168, 814–827 (2015).

19. Kim, D. Y., Bovet, L., Maeshima, M., Martinoia, E. & Lee, Y. The ABC transporter AtPDR8 is a cadmium extrusion pump conferring heavy metal resistance. Plant Journal 50, 207–218 (2007).

20. Takeuchi, M., Watanabe, A., Tamura, M. & Tsutsumi, Y. The gene expression analysis of Arabidopsis thaliana ABC transporters by real-time PCR for screening monolignol-transporter candidates. Journal of Wood Science 64, 477–484 (2018).

21. He, Y. et al. The Arabidopsis pleiotropic drug resistance transporters PEN3 and PDR12 mediate camalexin secretion for resistance to Botrytis cinerea. Plant Cell 31, 2206–2222 (2019).

22. Neelam, Khatkar, A. & Sharma, K. K. Phenylpropanoids and its derivatives: biological activities and its role in food, pharmaceutical and cosmetic industries. Critical Reviews in Food Science and Nutrition 60, 2655–2675 (2020).

23. Banasiak, J. et al. A Medicago truncatula ABC transporter belonging to subfamily G modulates the level of isoflavonoids. Journal of Experimental Botany 64, 1005–1015 (2013).

24. Biała, W., Banasiak, J., Jarzyniak, K., Pawela, A. & Jasiński, M. Medicago truncatula ABCG10 is a transporter of 4-coumarate and liquiritigenin in the medicarpin biosynthetic pathway. Journal of Experimental Botany 68, 3231–3241 (2017).

25. Orlando, B. J. & Liao, M. ABCG2 transports anticancer drugs via a closed-to-open switch. Nature Communications 11, 2264 (2020).

26. Kowal, J. et al. Structural basis of drug recognition by the multidrug transporter ABCG2. Journal of Molecular Biology 433, (2021).

27. Sun, Y. et al. Molecular basis of cholesterol efflux via ABCG subfamily transporters. Proceedings of the National Academy of Science USA 118, e2110483118 (2021).

28. Skarda, L., Kowal, J. & Locher, K. P. Structure of the human cholesterol transporter ABCG1. Journal of Molecular Biology 433, 167218 (2021).

29. Harris, A. et al. Structure and efflux mechanism of the yeast pleiotropic drug resistance transporter Pdr5. Nature Communications 12, 5254 (2021).

30. Khunweeraphong, N. & Kuchler, K. Multidrug resistance in mammals and fungi—from mdr to pdr: A rocky road from atomic structures to transport mechanisms. International Journal of Molecular Sciences 22, 4806 (2021).

31. Jumper, J. et al. Highly accurate protein structure prediction with AlphaFold. Nature 2021 596:7873 596, 583–589 (2021).

32. Hegedűs, T., Geisler, M., Lukács, G. L. & Farkas, B. Ins and outs of AlphaFold2 transmembrane protein structure predictions. Cellular and Molecular Life Sciences 79, 73 (2022).

33. Mariani, V., Biasini, M., Barbato, A. & Schwede, T. lDDT: A local superposition-free score for comparing protein structures and models using distance difference tests. Bioinformatics 29, 2722–2728 (2013).

34. Zhang, Y. & Skolnick, J. Scoring function for automated assessment of protein structure template quality. Proteins: Structure, Function, and Bioinformatics 57, 702–710 (2004).

35. Chovancova, E. et al. CAVER 3.0: A tool for the analysis of transport pathways in dynamic protein structures. PLOS Computational Biology 8, e1002708 (2012).

36. Brezovsky, J. et al. TransportTools: A library for high-throughput analyses of internal voids in biomolecules and ligand transport through them. Bioinformatics 38, 1752–1753 (2022).

37. Carpenter, E. J. et al. Access to RNA-sequencing data from 1,173 plant species: The 1000 Plant transcriptomes initiative (1KP). Gigascience 8, (2019).

38. Leebens-Mack, J. H. et al. One thousand plant transcriptomes and the phylogenomics of green plants. Nature 574, 679–685 (2019).

39. Ashkenazy, H. et al. ConSurf 2016: An improved methodology to estimate and visualize evolutionary conservation in macromolecules. Nucleic Acids Research 44, W344–W350 (2016).

40. Celniker, G. et al. ConSurf: Using evolutionary data to raise testable hypotheses about protein function. Israel Journal of Chemistry 53, 199–206 (2013).

41. Kamisetty, H., Ovchinnikov, S. & Baker, D. Assessing the utility of coevolution-based residue–residue contact predictions in a sequence- and structure-rich era. Proceedings of the National Academy of Science USA 110, 15674–15679 (2013).

42. Ovchinnikov, S., Kamisetty, H. & Baker, D. Robust and accurate prediction of residue-residue interactions across protein interfaces using evolutionary information. Elife 2014, (2014).

43. Jackson, S. M. et al. Structural basis of small-molecule inhibition of human multidrug transporter ABCG2. Nature Structural and Molecular Biology 25, 333–340 (2018).

44. Cunningham, B. C. & Wells, J. A. High-resolution epitope mapping of hGH-receptor interactions by alanine-scanning mutagenesis. Science 244, 1081–1085 (1989).

45. Toussaint, F., Pierman, B., Bertin, A., Lévy, D. & Boutry, M. Purification and biochemical characterization of NpABCG5/NpPDR5, a plant pleiotropic drug resistance transporter expressed in Nicotiana tabacum BY-2 suspension cells. Biochemical Journal 474, 1689–1703 (2017).

46. Vavra, O. et al. CaverDock: A molecular docking-based tool to analyse ligand transport through protein tunnels and channels. Bioinformatics 35, 4986–4993 (2019).

47. Papadopoulou, A. et al. Re-programming and optimization of a L-proline cis-4-hydroxylase for the cis-3-halogenation of its native substrate. ChemCatChem 13, 3914–3919 (2021).

48. Marques, S. M., Bednar, D. & Damborsky, J. Computational study of protein-ligand unbinding for enzyme engineering. Frontiers in Chemistry 7, 650 (2019).

49. Brodsky, K. et al. Dual substrate specificity of the rutinosidase from Aspergillus niger and the role of its substrate tunnel. International Journal of Molecular Sciences 21, 5671 (2020).

50. Thomas, C. & Tampé, R. Structural and mechanistic principles of ABC transporters. Annual Review of Biochemistry 89, 605–636 (2020).

51. Khan, A. et al. Antifungal activity of plant secondary metabolites on Candida albicans: An updated review. Current Molecular Pharmacology 15, (2022).

52. Moreno, A., Banerjee, A., Prasad, R. & Falson, P. PDR-like ABC systems in pathogenic fungi. Research in Microbiology 170, 417–425 (2019).

53. Nagata, T., Nemoto, Y. & Hasezawa, S. Tobacco BY-2 cell line as the “HeLa” cell in the cell biology of higher plants. International Review of Cytology 132, 1–30 (1992).

54. Curtis, M. D. & Grossniklaus, U. A Gateway cloning vector set for high-throughput functional analysis of genes in planta. Plant Physiology 133, 462–469 (2003).

55. Hellens, R., Mullineaux, P. & Klee, H. A guide to Agrobacterium binary Ti vectors. Trends in Plant Science 5, 446–451 (2000).

56. Jasiński, M. et al. A plant plasma membrane ATP binding cassette-type transporter is involved in antifungal terpenoid secretion. Plant Cell 13, 1095–107 (2001).

57. Larsson, C., Widell, S. & Kjellbom, P. [52] Preparation of high-purity plasma membranes. in Methods in Enzymology 148, 558–568 (1987).

58. Edgar, R. C. MUSCLE: Multiple sequence alignment with high accuracy and high throughput. Nucleic Acids Research 32, 1792–1797 (2004).

59. Kumar, S., Stecher, G., Li, M., Knyaz, C. & Tamura, K. MEGA X: Molecular evolutionary genetics analysis across computing platforms. Molecular Biology and Evolution 35, 1547–1549 (2018).

60. Laskowski, R. A., MacArthur, M. W., Moss, D. S. & Thornton, J. M. PROCHECK: A program to check the stereochemical quality of protein structures. Journal of Applied Crystallography 26, 283–291 (1993).

61. Trott, O. & Olson, A. J. AutoDock Vina: Improving the speed and accuracy of docking with a new scoring function, efficient optimization, and multithreading. Journal of Computational Chemistry 31, 455–461 (2010).

62. D.A. Case et al. AMBER 20. Preprint at http://ambermd.org/ (2020).

63. Anandakrishnan, R., Aguilar, B. & Onufriev, A. V. H++ 3.0: automating pK prediction and the preparation of biomolecular structures for atomistic molecular modeling and simulations. Nucleic Acids Research 40, W537–W541 (2012).

64. Jo, S., Kim, T., Iyer, V. G. & Im, W. CHARMM-GUI: A web-based graphical user interface for CHARMM. Journal of Computational Chemistry 29, 1859–1865 (2008).

65. Brooks, B. R. et al. CHARMM: The biomolecular simulation program. Journal of Computational Chemistry 30, 1545–1614 (2009).

66. Lee, J. et al. CHARMM-GUI input generator for NAMD, GROMACS, AMBER, OpenMM, and CHARMM/OpenMM simulations using the CHARMM36 additive force field. Journal of Chemical Theory and Computation 12, 405–413 (2016).

67. Lee, J. et al. CHARMM-GUI supports the Amber force fields. The Journal of Chemical Physics 153, 035103 (2020).

68. Wu, E. L. et al. CHARMM-GUI Membrane Builder toward realistic biological membrane simulations. Journal of Computational Chemistry 35, 1997–2004 (2014).

69. Izadi, S., Anandakrishnan, R. & Onufriev, A. V. Building water models: A different approach. Journal of Physical Chemistry Letters 5, 3863–3871 (2014).

70. Hopkins, C. W., Le Grand, S., Walker, R. C. & Roitberg, A. E. Long-time-step molecular dynamics through hydrogen mass repartitioning. Journal of Chemical Theory and Computation 11, 1864–1874 (2015).

71. Le Grand, S., Götz, A. W. & Walker, R. C. SPFP: Speed without compromise—A mixed precision model for GPU accelerated molecular dynamics simulations. Computer Physics Communications 184, 374–380 (2013).

72. Tian, C. et al. Ff19SB: Amino-acid-specific protein backbone parameters trained against quantum mechanics energy surfaces in solution. Journal of Chemical Theory and Computation 16, 528–552 (2020).

73. Meagher, K. L., Redman, L. T. & Carlson, H. A. Development of polyphosphate parameters for use with the AMBER force field. Journal of Computational Chemistry 24, 1016–1025 (2003).

74. Morris, G. M. et al. AutoDock4 and AutoDockTools4: Automated docking with selective receptor flexibility. Journal of Computational Chemistry 30, 2785–2791 (2009).

75. Roe, D. R. & Cheatham, T. E. PTRAJ and CPPTRAJ: Software for processing and analysis of molecular dynamics trajectory data. Journal of Chemical Theory and Computation 9, 3084–3095 (2013).

76. Michaud-Agrawal, N., Denning, E. J., Woolf, T. B. & Beckstein, O. MDAnalysis: A toolkit for the analysis of molecular dynamics simulations. Journal of Computational Chemistry 32, 2319–2327 (2011).

77. Gowers, R. J. et al. MDAnalysis: A Python package for the rapid analysis of molecular dynamics simulations. Proceedings of the 15th Python in Science Conference 98–105 (2016) doi:10.25080/MAJORA-629E541A-00E.

