## Supplementary Figures and Tables for "Restriction of access to the central cavity is a major contributor to substrate selectivity in plant ABCG transporters"

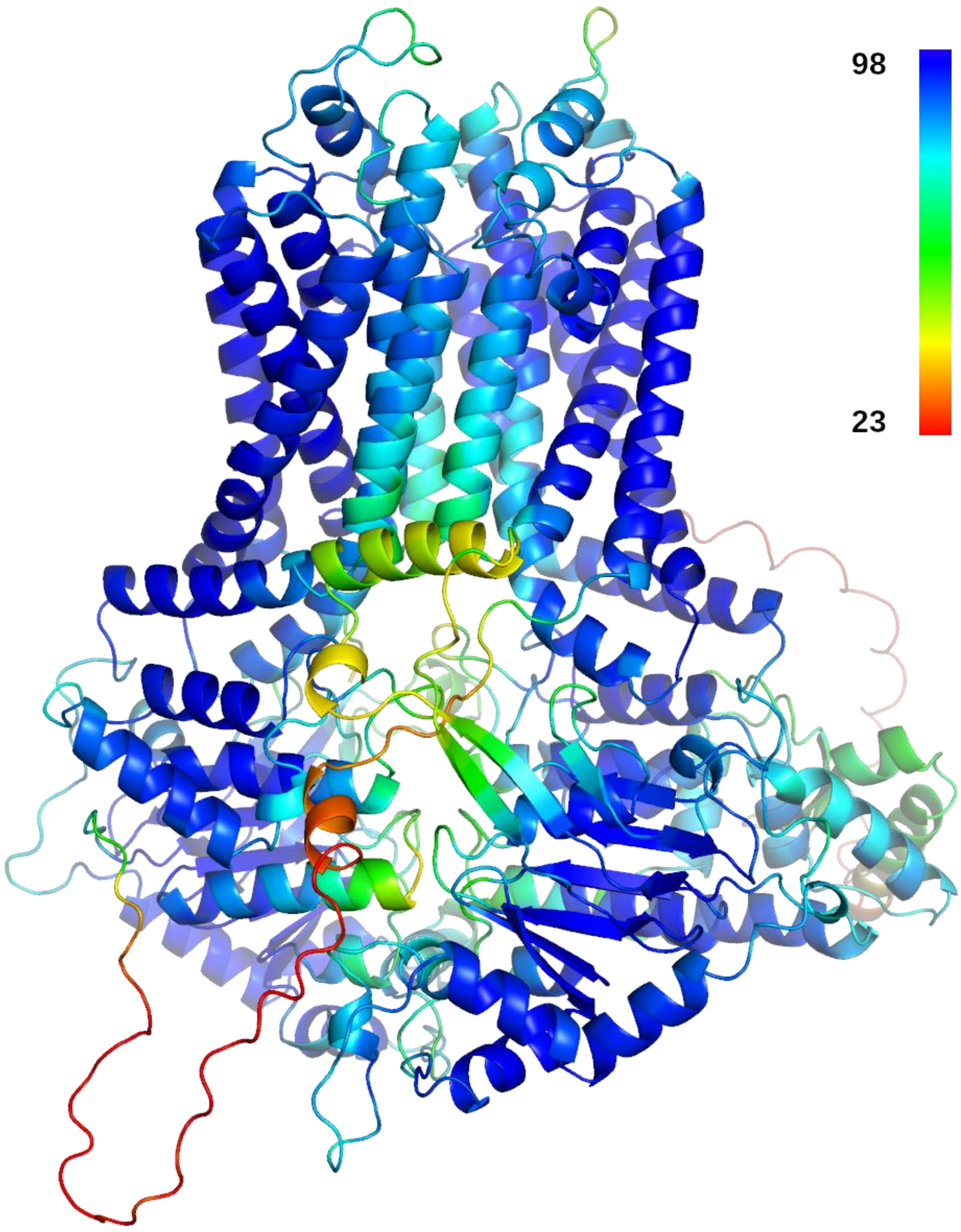


**Supplementary Figure 1: AlphaFold2 model of MtABCG46.** Protein model colored by per-residue lDDT-Cα score. Regions with low confidence are colored in red, orange, and yellow.

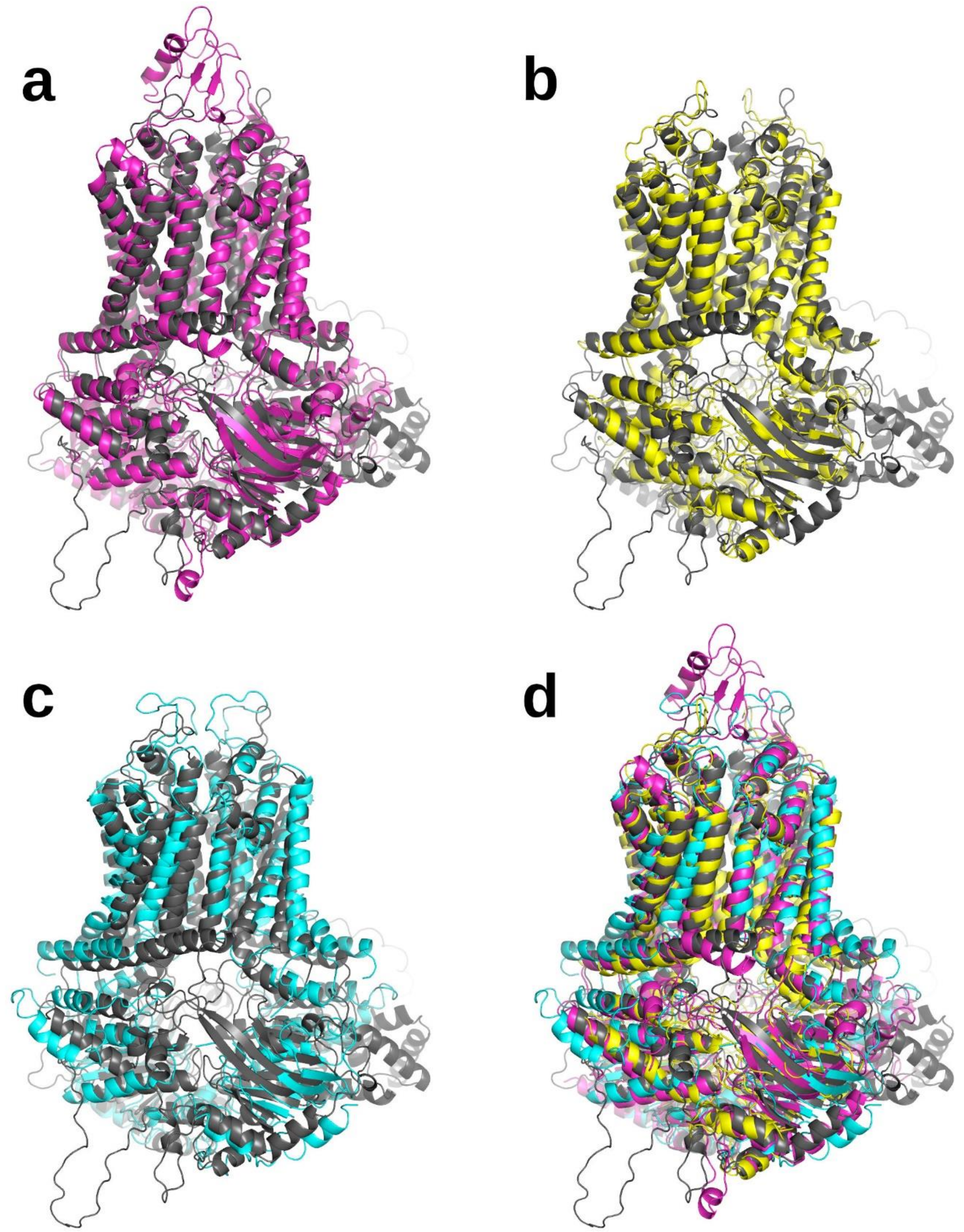


**Supplementary Figure 2: Structural alignment of MtABCG46 related ABCG transporters. a-d** Structural alignment to MtABCG46 in gray obtained with TM-score for ABCG related proteins: ScPDR5 in magenta (PDB id: 7p04) (**a**); HsABCG1 in yellow (PDB id: 7r8e) (**b**); HsABCG2 in cyan (PDB id: 6vxh) (**c**); all four structures aligned show the same fold (**d**).

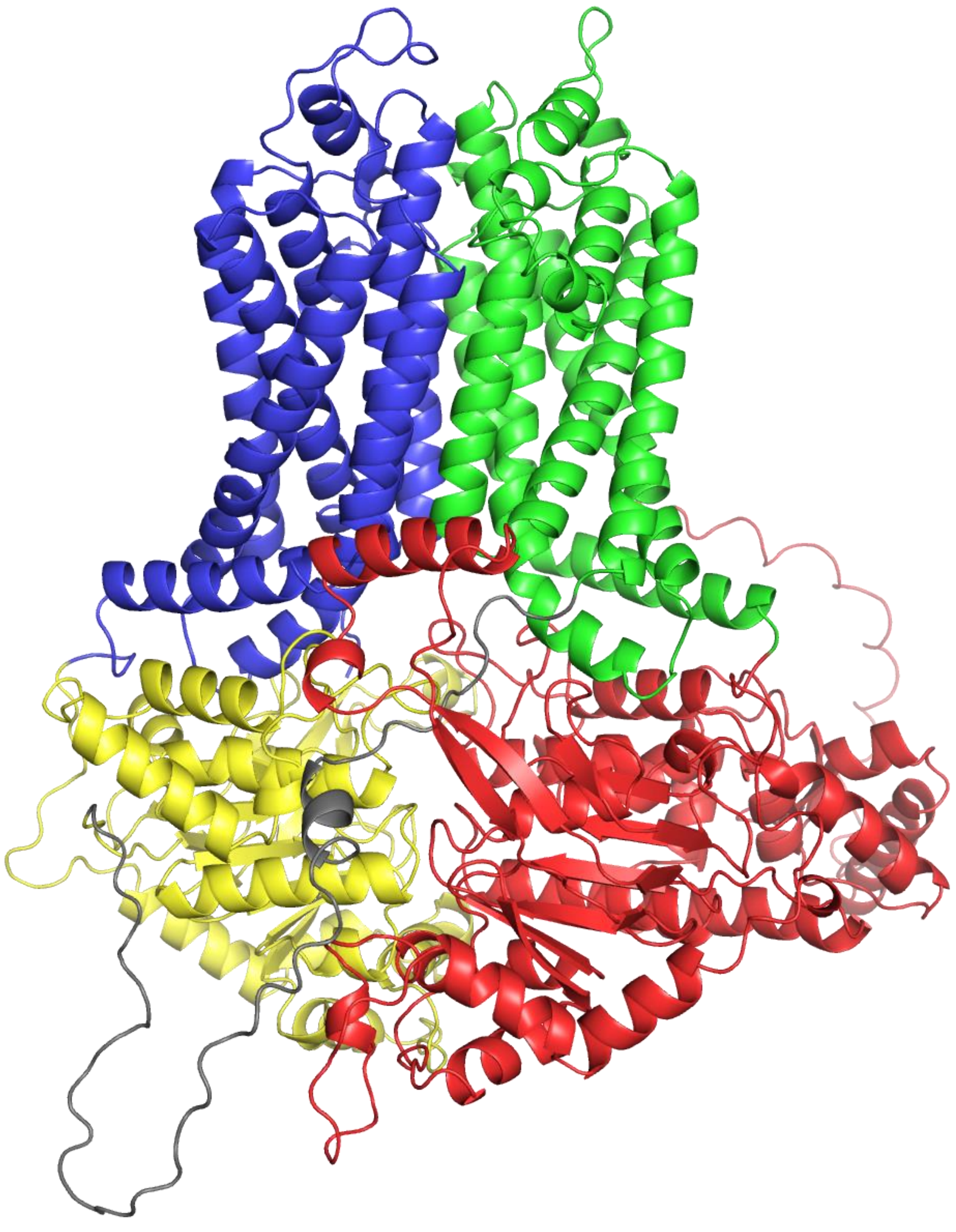

**Supplementary Figure 3: Domains typical for full-size ABCG structures present in MtABCG46.** NBD1 in red, TMD1 in green, linker region in gray, NBD2 in yellow, and TMD2 in blue.

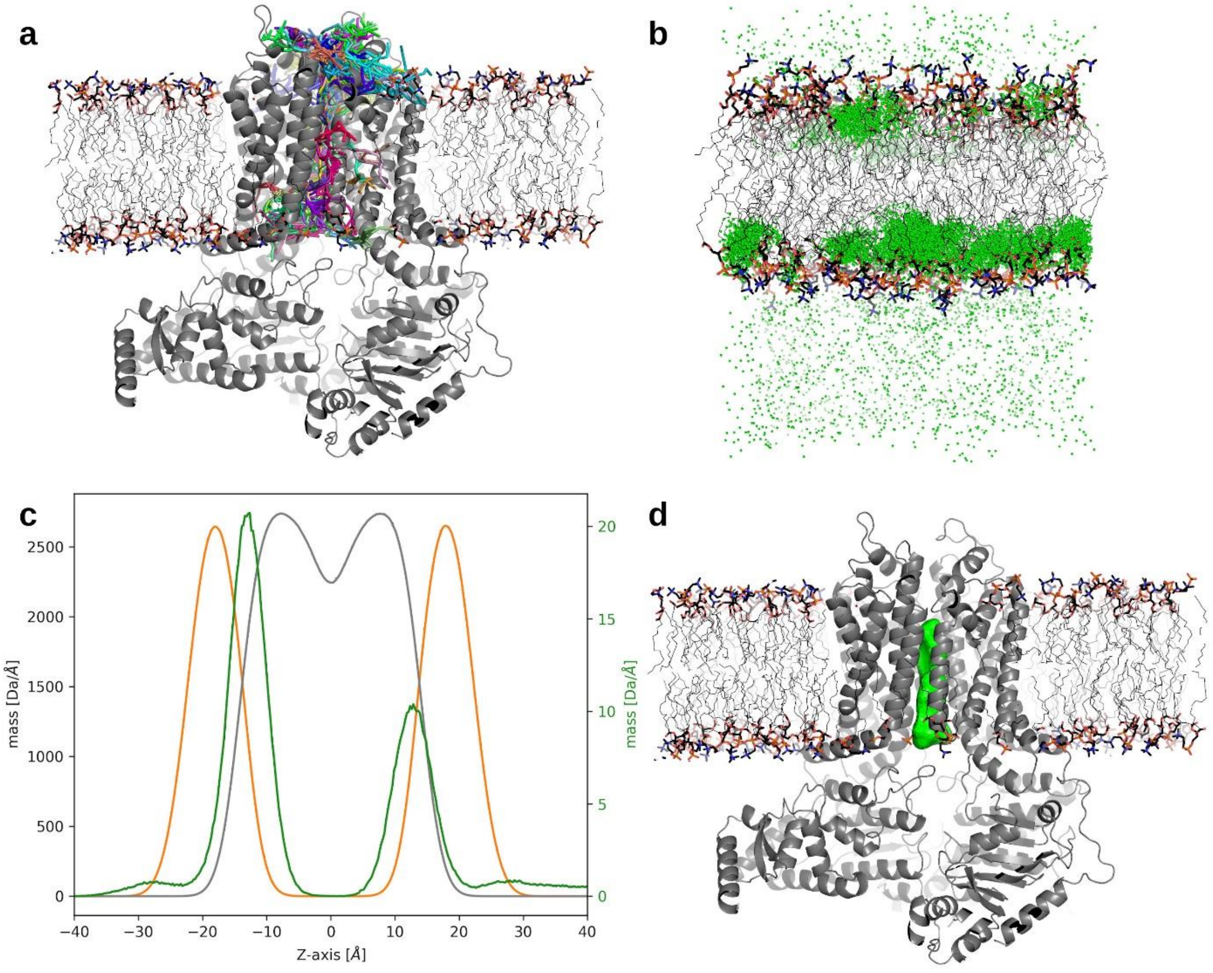


**Supplementary Figure 4: Network of access paths in MtABCG46 obtained with TransportTools and liquiritigenin localization. a** Clusters of paths (differentially colored sticks) identified in MtABCG46 structure (gray cartoon) embedded in POPC membrane shown with heads as sticks and tails as lines. **b** Preferred liquiritigenin localization in the membrane; each green sphere represents the center of mass of a liquiritigenin molecule per frame. **c** Linear mass density across the Z-axis, POPC heads colored orange and POPC tails as gray with left axis; liquiritigenin colored as green with the right axis. **d** MtABCG46 structure as a gray cartoon with the overall volume of selected access path ensemble represented as a green surface.

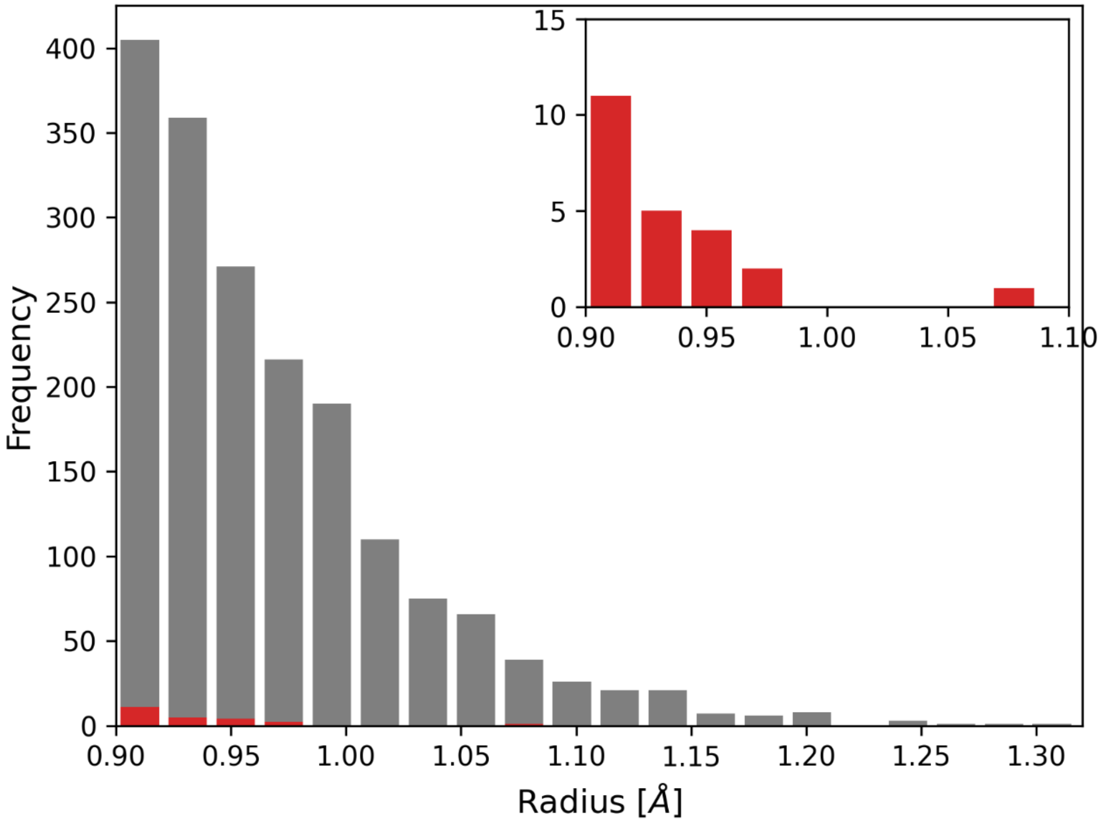


**Supplementary Figure 5: Presence of the access path in MtABCG46 ATP-bound and APO (without ATP and $MG^{2+}$ ions) models.** The amount of frames where the access path was present in the ATP-bound model is colored in gray and in red for the APO form. In the inset, only the results from the APO variant are shown.

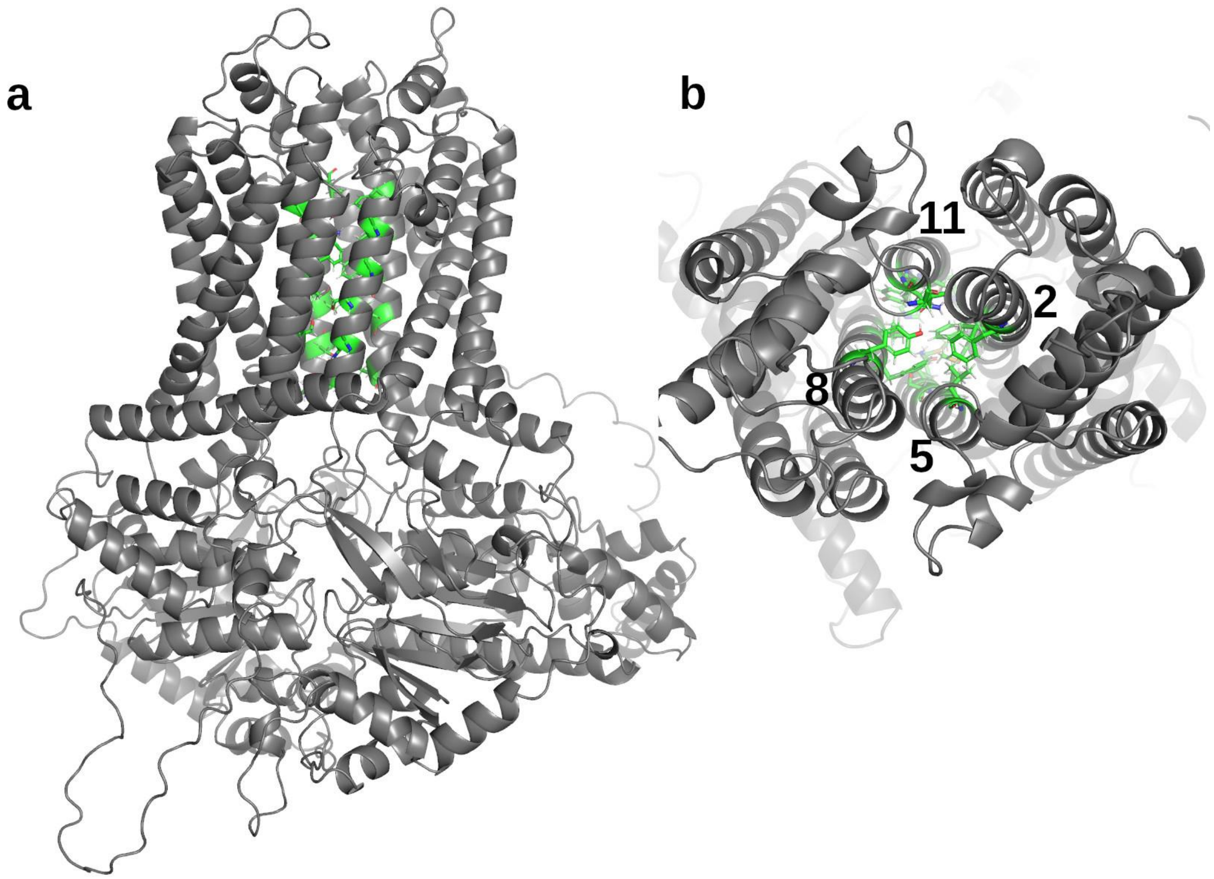


**Supplementary Figure 6: Space between four TMD helices harboring the central cavity and its access path in MtABCG46. a** Side view of the model showing four TMD helices. **b** Top view of the lining residues.

**a**

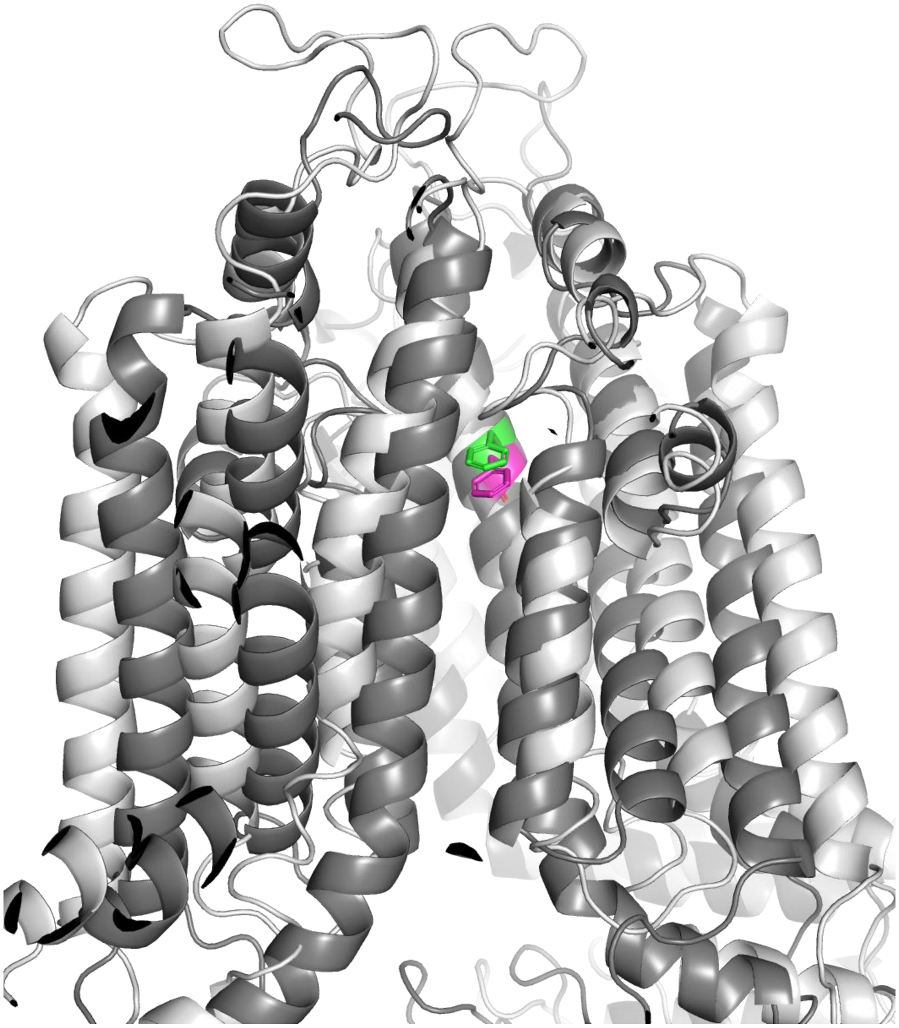

**b**

```
MtABCG46     532   IMALIAMTLFFRTEMHRNNQDDAGVY--AGALFFTLVTMMFNGMSEISMT  579
                   ++ L+   ++F  +      D  G+   AG LFF      F+ +S + +
HsABCG2      404   VLGLVIGAIYFGLK-----NDSTGIQNRAGVLFFLTTNQCFSSVSAVELF  448
```

**Supplementary Figure 7: Structural and sequential alignment of MtABCG46 and HsABCG2. a** Structure of HsABCG2 in a light gray cartoon with residue F431 highlighted as magenta sticks, and MtABCG46 as a dark gray cartoon with residue highlighted F562 as green sticks. **b** Needleman-Wunsch sequence alignment of MtABCG46 and HsABCG2 dimer. F562 of MtABCG46 and corresponding residue of HsABCG2 – F431 marked bold and yellow.

| TAXA | F562 | M573 | S577 | M664 | N668 | V675 | L679 | V1221 | S1225 | H1316 | H1317 | S1320 |
|---|---|---|---|---|---|---|---|---|---|---|---|---|
| Green Algae | F(s) | F | A | M | N | L | F | M | S | Q | Q | A |
| Bryophytes | F(s) | F | S | M | N | L | F | V | S | H | Q | A |
| Pteridophytes | F(s) | F | S | M | N | L | F | I | S | H | Q | A |
| Monocots | F | F | A | M | N | L | L | I | S | Y | Q/N | A |
| Core eudicots | F | F | S | M | N | L | F | I | S | H | Q | A |

| Conservation scale | | | | | | | | |
|---|---|---|---|---|---|---|---|---|
| 1 | 2 | 3 | 4 | 5 | 6 | 7 | 8 | 9 |
| Variable | | | | Average | | | | Highly conserved |

**Supplementary Figure 8: Conservation score of the inner cavity forming residues among different plant taxa based on Consurf analysis.** The most commonly present amino acid in the corresponding residue within taxa is indicated by a single letter amino acid code; (s) corresponds to residues predicted by Consurf as structural.

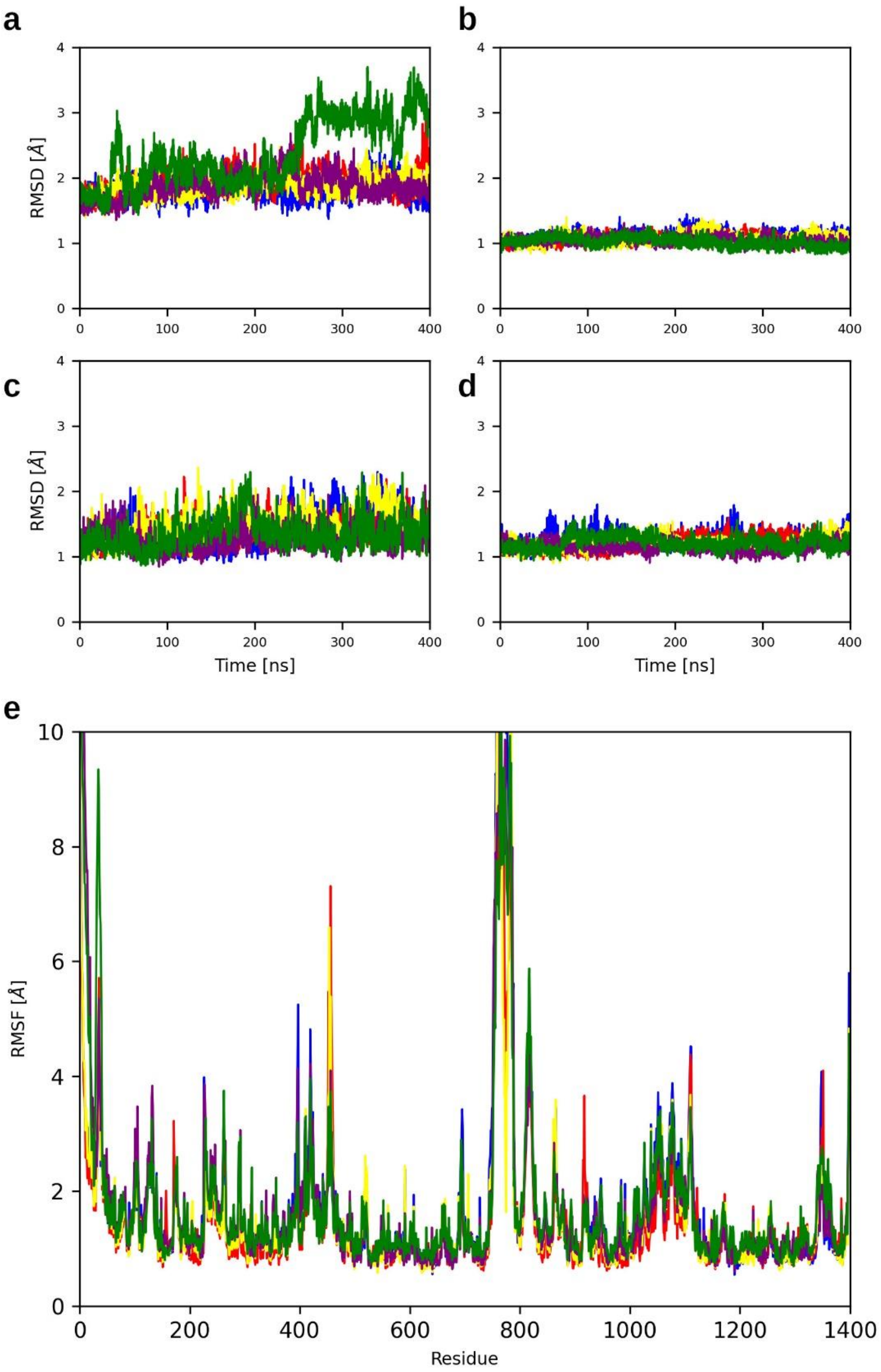


**Supplementary Figure 9: RMSD and RMSF of five independent replicas of MtABCG46. a-e** RMSD plots separated by regions: NBD1 (**a**), TMD1 (**b**), NBD2 (**c**) and TMD2 (**d**) and RMSF (**e**).

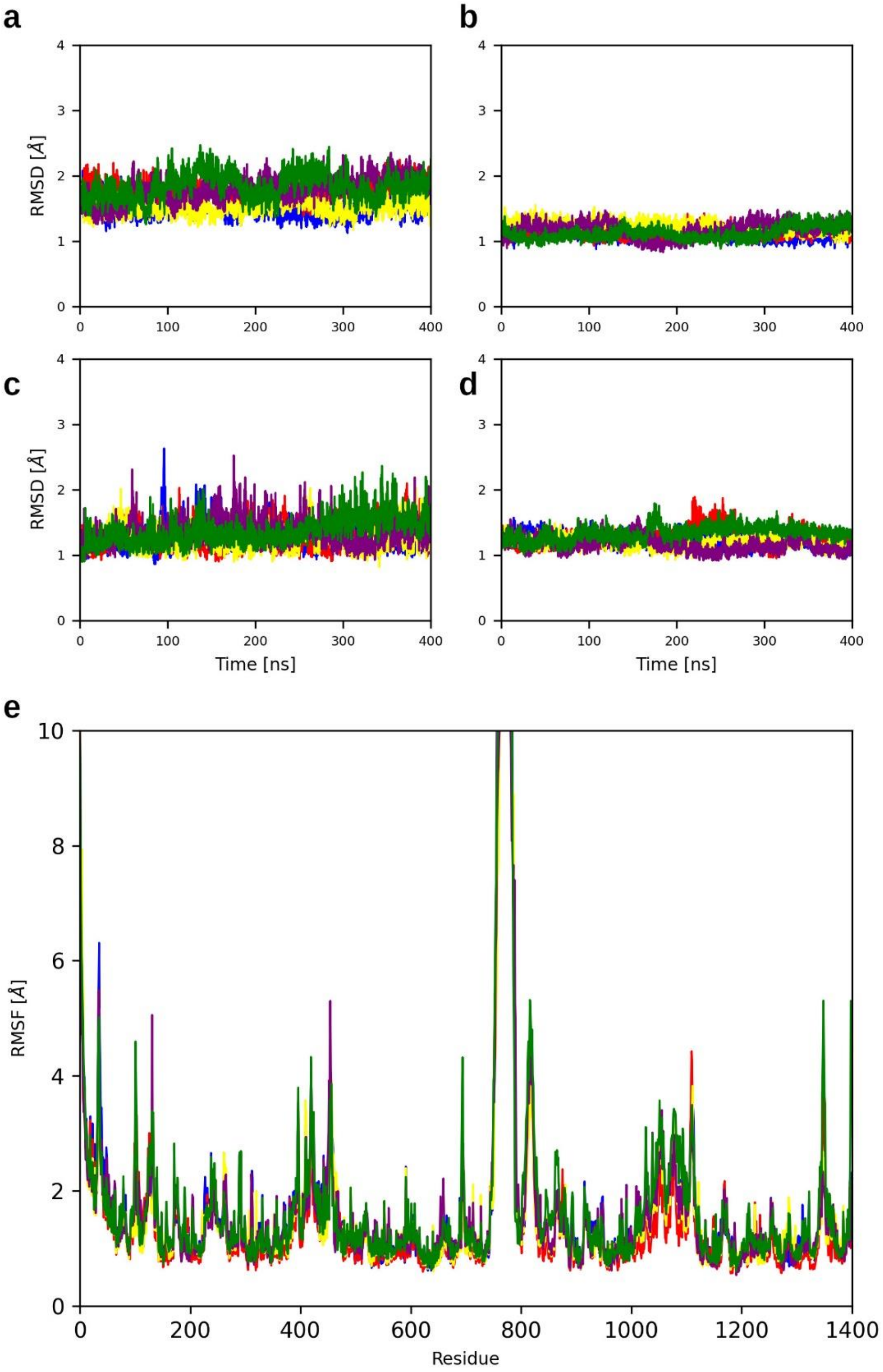


**Supplementary Figure 10: RMSD and RMSF of five independent replicas of MtABCG46 F562L mutant. a-e** RMSD plots separated by regions: NBD1 (**a**), TMD1 (**b**), NBD2 (**c**) and TMD2 (**d**) and RMSF (**e**).

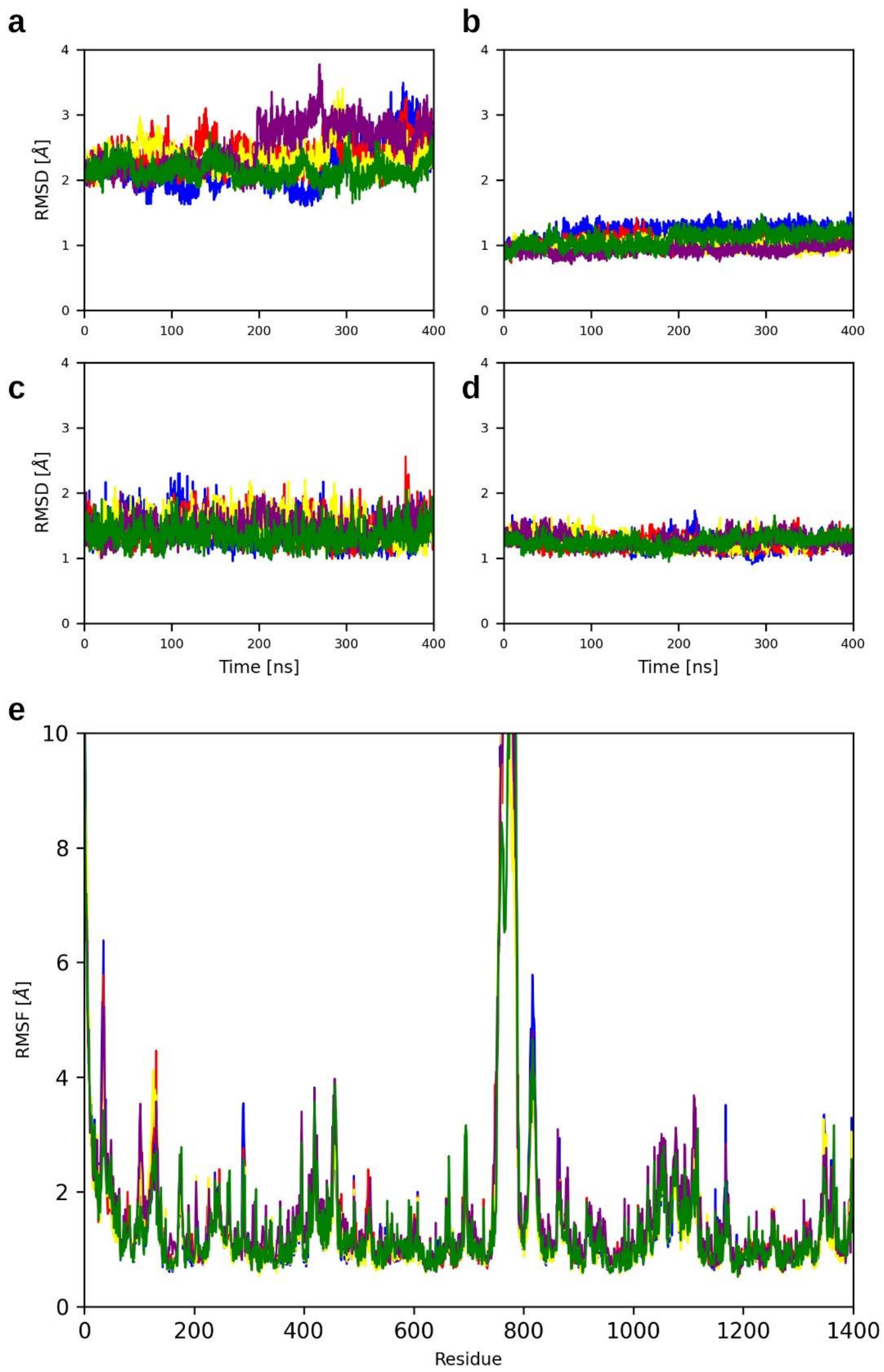


**Supplementary Figure 11: RMSD and RMSF of five independent replicas of MtABCG46 F562Y mutant. a-e** RMSD plots separated by regions: NBD1 (**a**), TMD1 (**b**), NBD2 (**c**) and TMD2 (**d**) and RMSF (**e**).

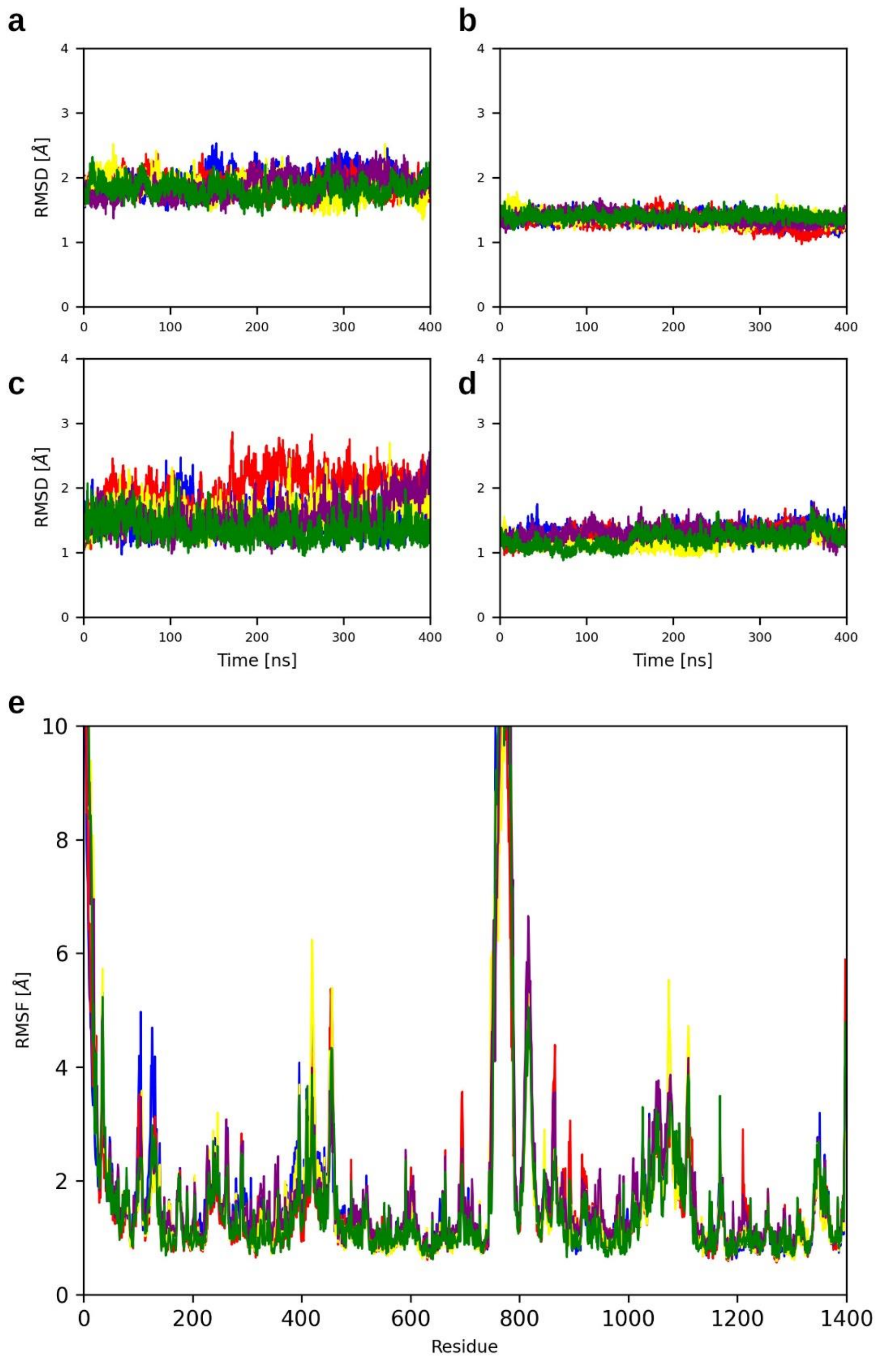


**Supplementary Figure 12: RMSD and RMSF of five independent replicas of MtABCG46 F562A mutant. a-e** RMSD plots separated by regions: NBD1 (**a**), TMD1 (**b**), NBD2 (**c**) and TMD2 (**d**) and RMSF (**e**).

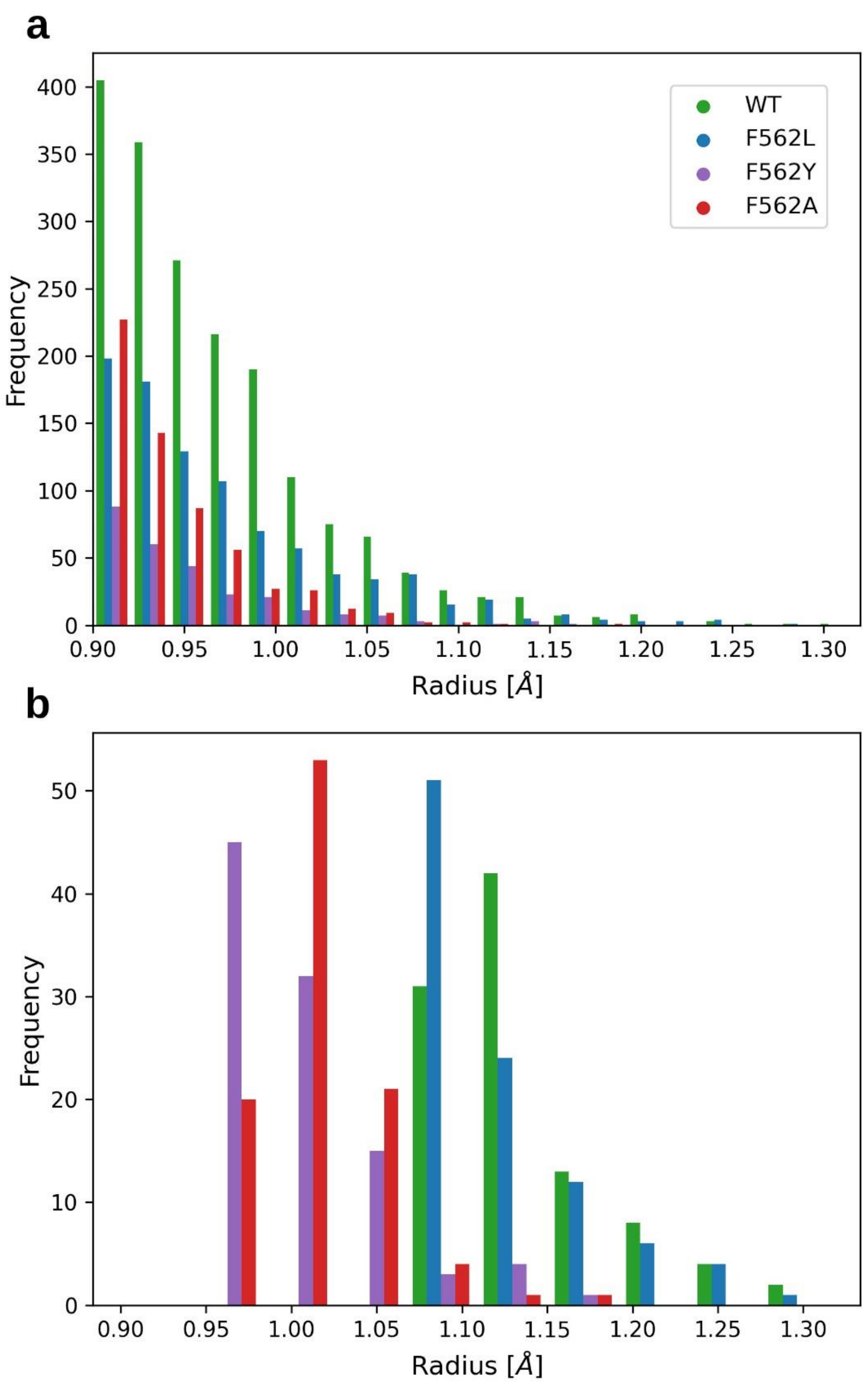


**Supplementary Figure 13: Population of access paths by their bottleneck radius. a, b** The access path population is shown for all the paths present in each variant (**a**), the 100 widest paths (**b**).

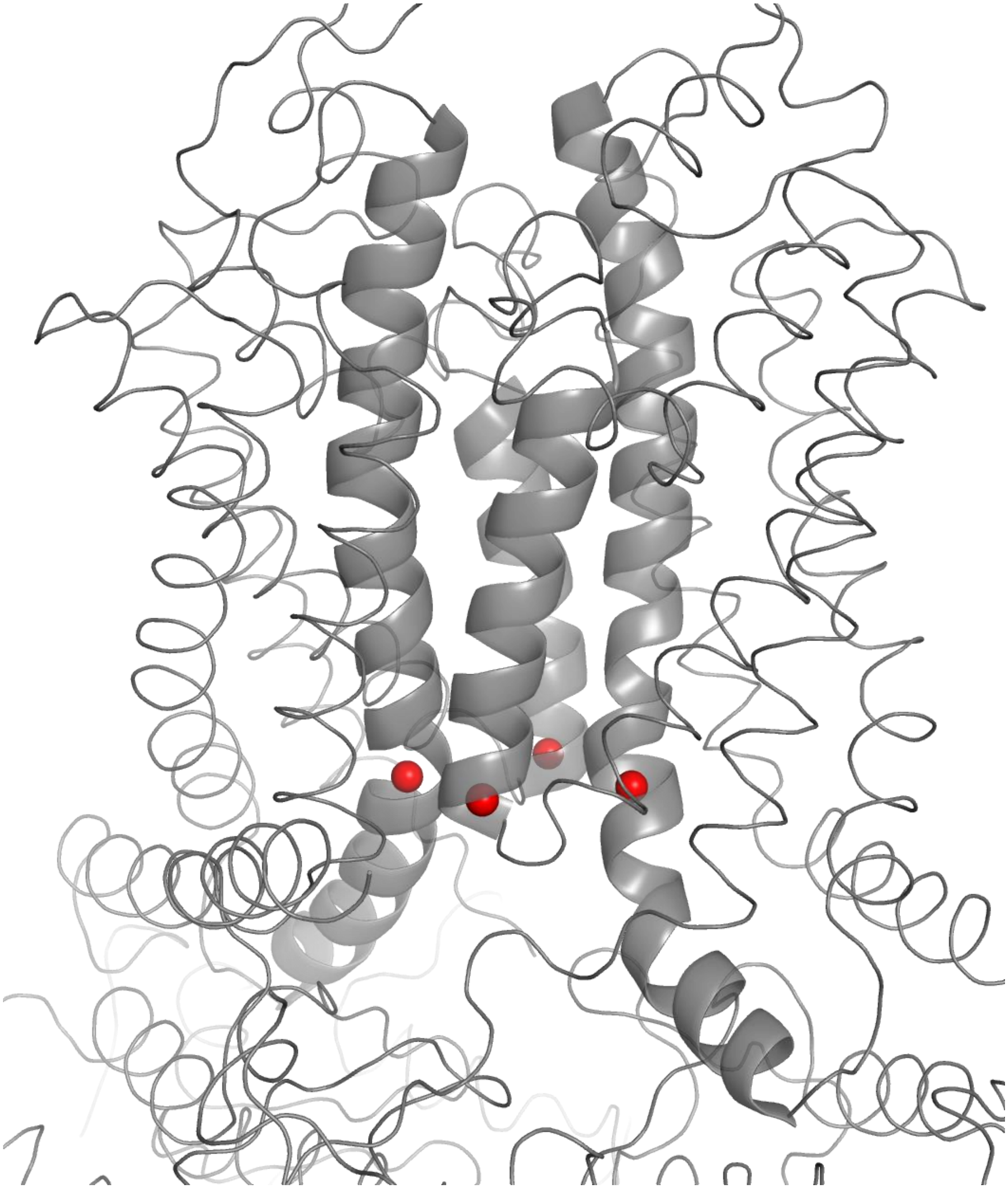

**Supplementary Figure 14: Area of the entrance to the access path in MtABCG46.** The four TMD helices 2, 5, 8, and 11 are represented as a cartoon, with the entry region delimited by the four red spheres corresponding to the helix centerlines defined by the HELANAL module of MDAnalysis.

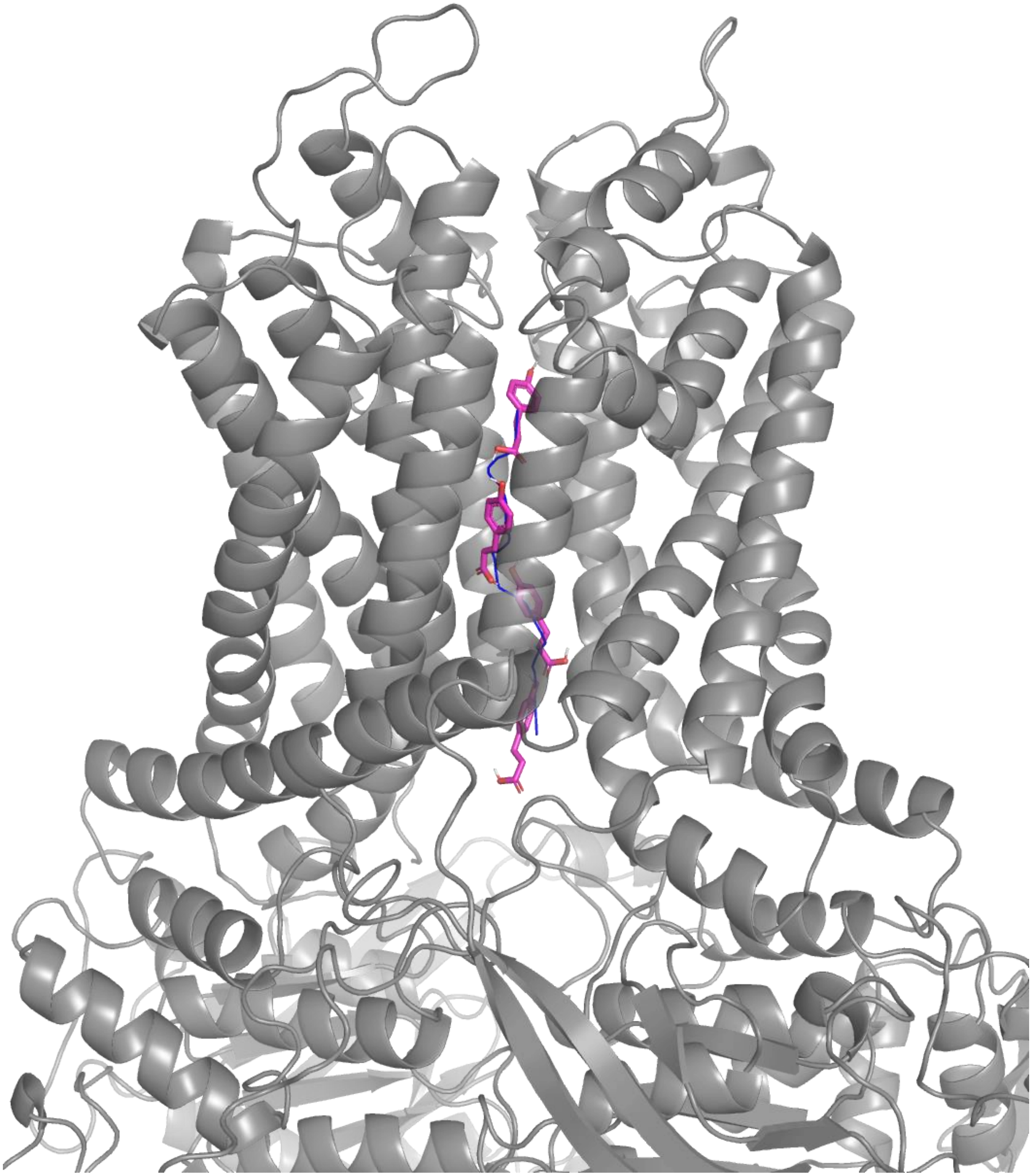

**Supplementary Figure 15: Migration from the intracellular region to the internal cavity in MtABCG46.** An example of the migration of 4-coumarate (magenta sticks) across the access path in MtABCG46, four snapshots are depicted along the path centerline (blue stick).

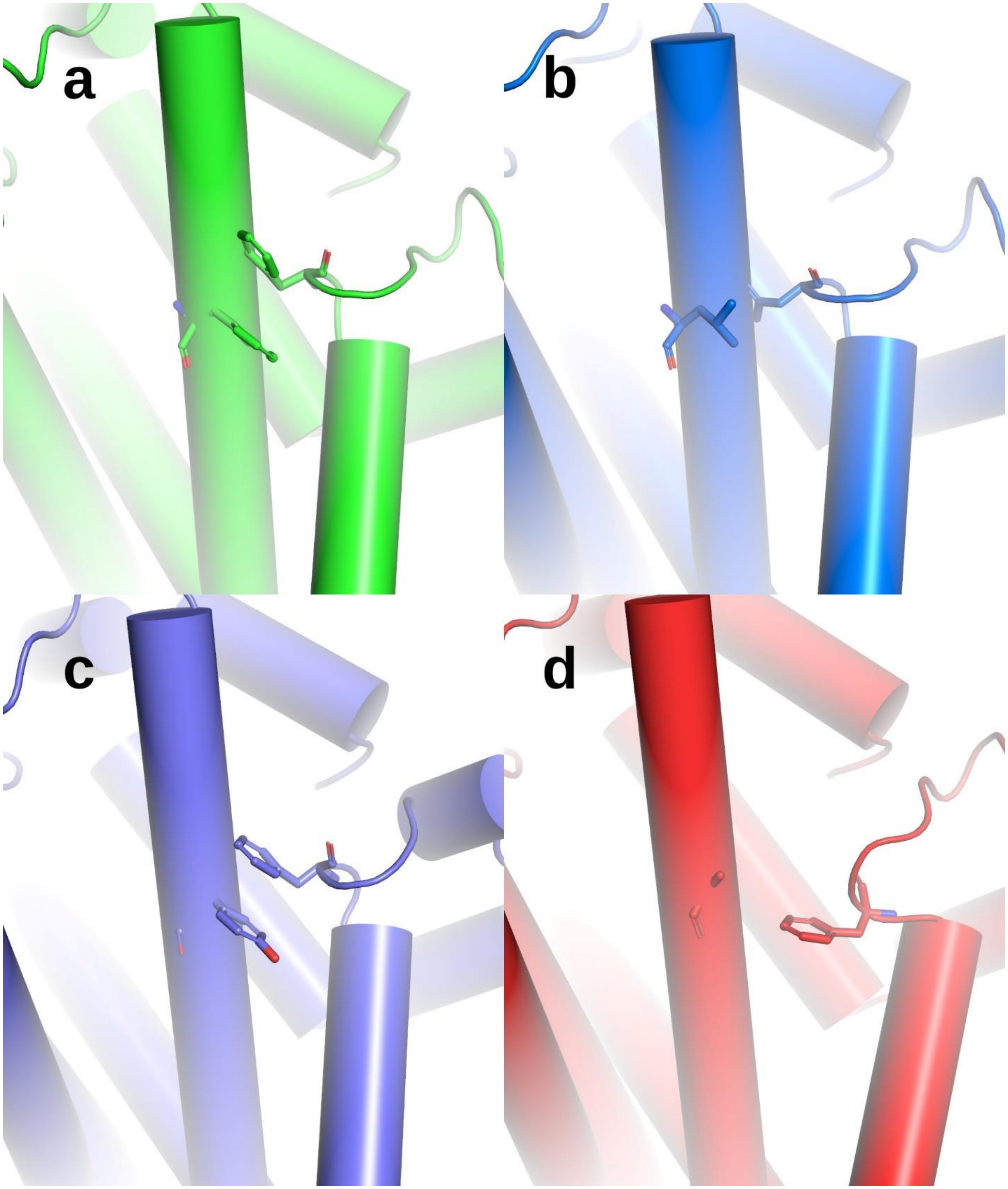


**Supplementary Figure 16: Parallel displaced π-stacking interaction between F684 and altered residues at position 562. a-d** Sliced TM lateral view of MtABCG46, showcasing the non-covalent contacts between F684 and altered residues (in the sticks) as follows: F562 in WT (**a**), L562 in leucine mutant (**b**), Y562 in tyrosine mutant (**c**) and A562 in alanine mutant (**d**).

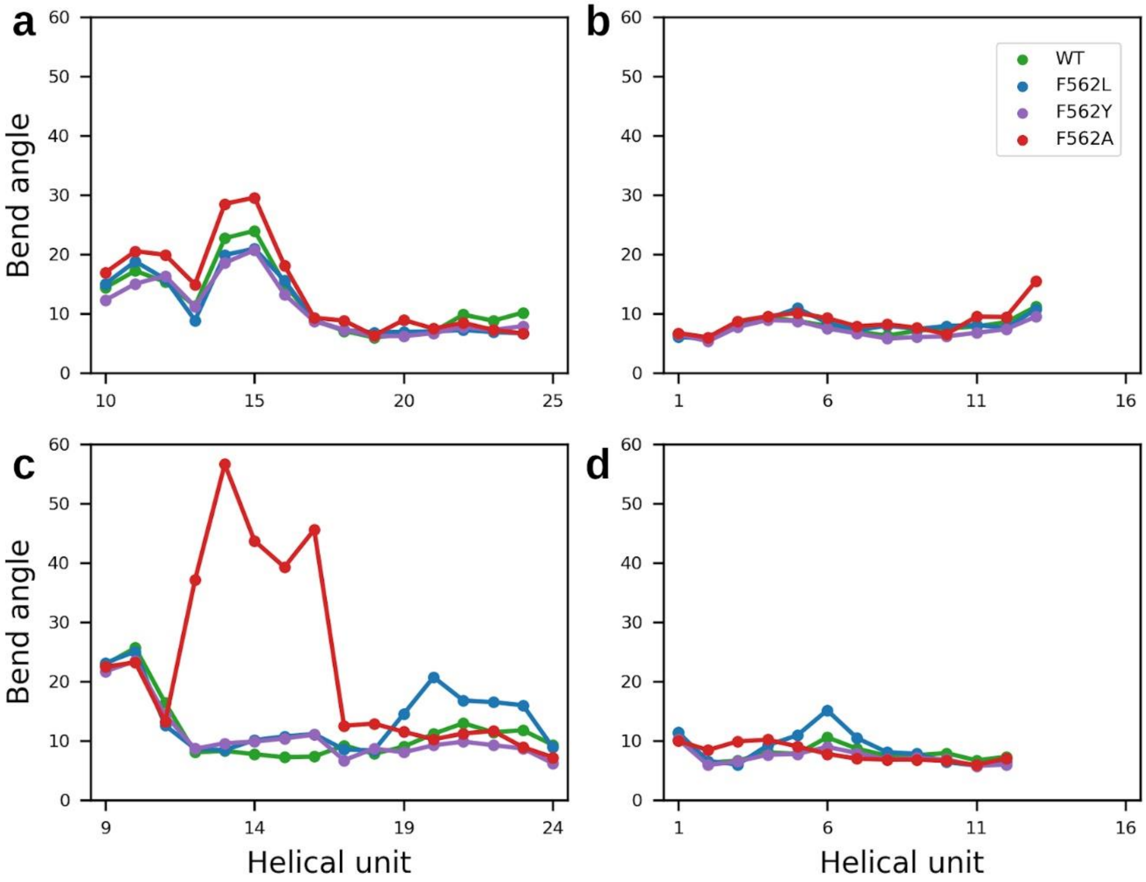


**Supplementary Figure 17: Bending angles of TMD helices forming the access path in MtABCG46 WT and mutants.** The angles (in degrees) for helical units defined by the HELANAL module of MDAnalysis are presented for: **a** TMD helix 2, **b** TMD helix 5, **c** TMD helix 8, and **d** TMD helix 11. Only the regions covering the entrance path are shown. The averages for five replications of each variant are shown as dots.

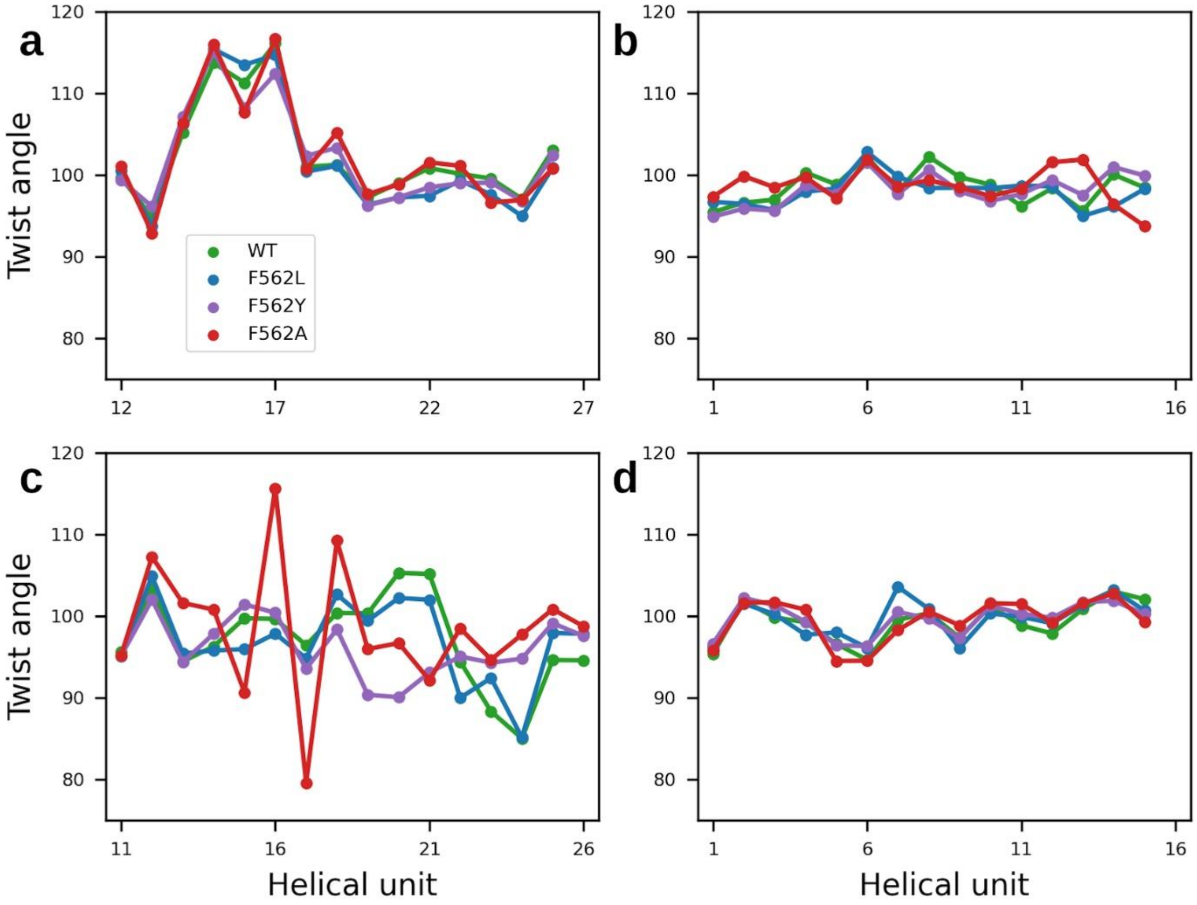


**Supplementary Figure 18: Twisting angles of TMD helices forming the access path in MtABCG46 WT and mutants.** The angles (in degrees) for helical units defined by the HELANAL module of MDAnalysis are presented for: **a** TMD helix 2, **b** TMD helix 5, **c** TMD helix 8, and **d** TMD helix 11. Only the regions covering the entrance path are shown. The averages for five replications of each variant are shown as dots.

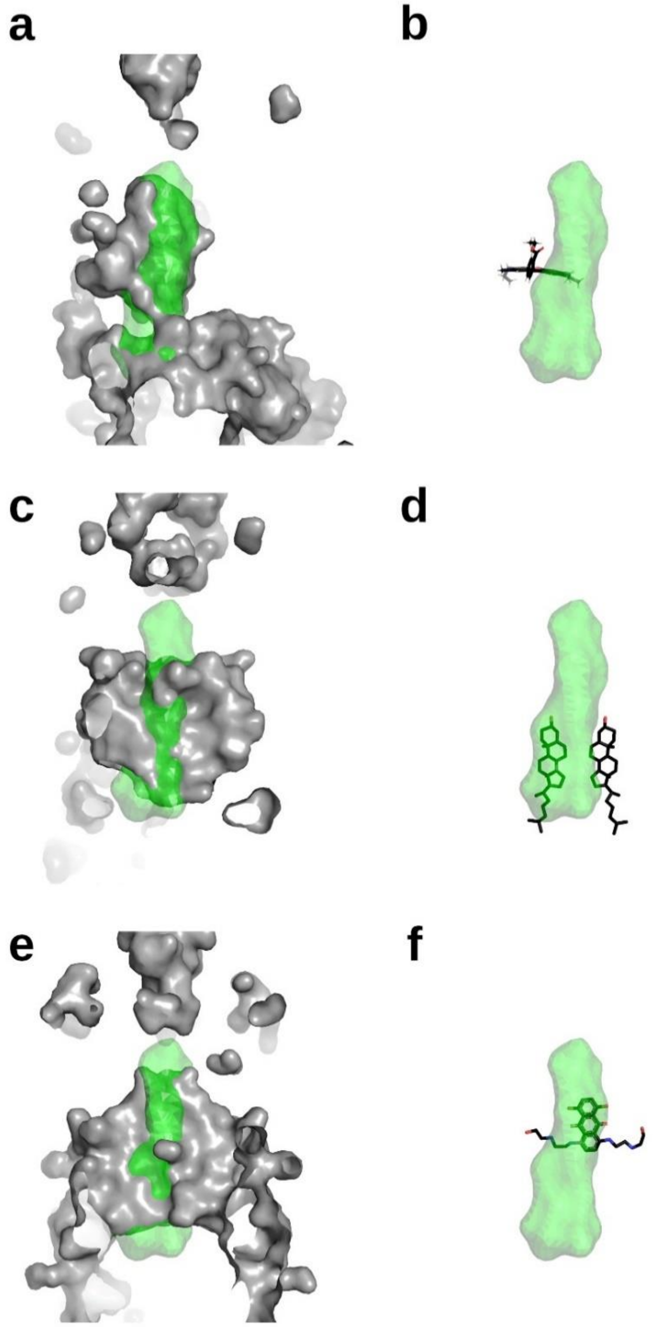


**Supplementary Figure 19: Cavities in related ABCG transporters and presence of ligands therein. a, c, e** Comparison between the access path ensemble in MtABCG46 shown as green surface and cavities found in experimental structures of related ABCG transporters shown as light gray surfaces: ScPDR5 (**a**), HsABCG1 (**c**), HsABCG2 (**e**). **b**, **d**, **f** Ligands present in related ABCG transporters superposed on the volume of the access path in MtABCG46 are shown as sticks: rhodamine 6G in ScPDR5 (**b**), cholesterol in HsABCG1 (**d**) mitoxantrone in HsABCG2 (**f**). All the structures were aligned with TM-align.

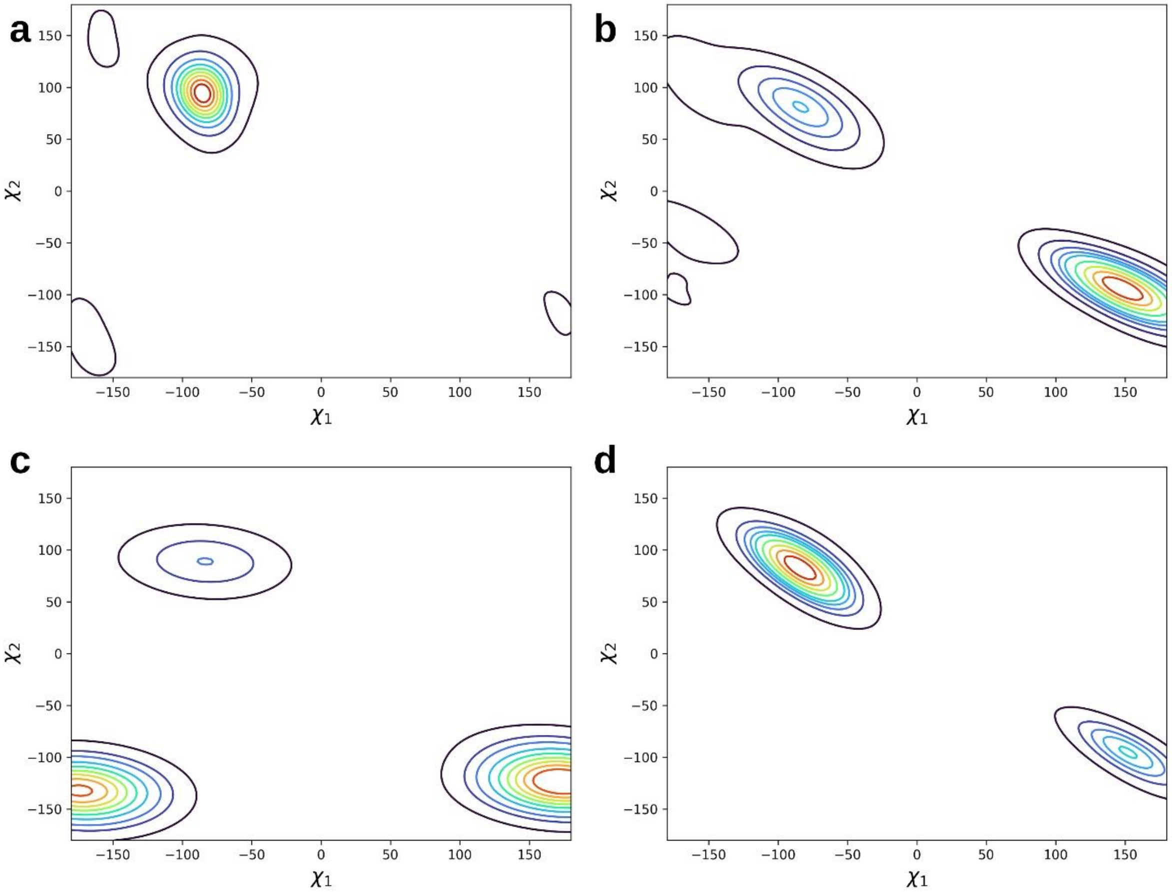


**Supplementary Figure 20: Conformational space adopted by N1331. a-d** Kernel density estimate plots for the chi angles of N1331 in MtABCG46 variants WT (**a**), F562L (**b**), F562Y (**c**) and F562A (**d**) mutants.

**Supplementary Table 1. Top ten access path ensembles identified with TransportTools from five simulations of MtABCG46**

| ID | MD runs | Total Frames | Avg Frames | Avg BR ±StDev | Max BR | Avg Length ±StDev | Avg Curv ±StDev | Avg Throughput ±StDev | Priority | Exit region | Frequency |
|---|---|---|---|---|---|---|---|---|---|---|---|
| 1 | 4 | 2290 | 458 | 1.10 ±0.19 | 2.06 | 24.76 ±2.91 | 1.27 ±0.14 | 0.49 ±0.09 | 0.223 | EC | 11.45% |
| 2 | 5 | 1402 | 280.4 | 1.03 ±0.13 | 1.72 | 28.90 ±4.75 | 1.39 ±0.13 | 0.39 ±0.08 | 0.135 | EC | 7.01% |
| **3** | **5** | **1816** | **363.2** | **0.97 ±0.06** | **1.32** | **34.58 ±2.94** | **1.33 ±0.09** | **0.19 ±0.04** | **0.087** | **IC** | **9.08%** |
| 4 | 3 | 1540 | 308 | 1.09 ±0.17 | 1.9 | 32.27 ±4.88 | 1.63 ±0.19 | 0.38 ±0.08 | 0.087 | EC | 7.70% |
| 5 | 4 | 938 | 187.6 | 1.07 ±0.15 | 1.7 | 26.82 ±5.32 | 1.36 ±0.22 | 0.39 ±0.10 | 0.073 | EC | 4.69% |
| 6 | 4 | 754 | 150.8 | 1.03 ±0.13 | 1.65 | 27.41 ±4.28 | 1.28 ±0.12 | 0.40 ±0.09 | 0.061 | EC | 3.77% |
| 7 | 3 | 995 | 199 | 1.07 ±0.16 | 1.71 | 35.45 ±4.55 | 1.71 ±0.17 | 0.30 ±0.07 | 0.045 | EC | 4.98% |
| 8 | 5 | 637 | 127.4 | 0.97 ±0.06 | 1.26 | 38.26 ±4.11 | 1.44 ±0.17 | 0.17 ±0.05 | 0.026 | IC | 3.19% |
| 9 | 2 | 734 | 146.8 | 1.04 ±0.14 | 1.55 | 30.46 ±4.43 | 1.57 ±0.18 | 0.35 ±0.08 | 0.025 | EC | 3.67% |
| 10 | 1 | 851 | 170.2 | 1.11 ±0.18 | 2.12 | 25.92 ±3.26 | 1.24 ±0.10 | 0.48 ±0.09 | 0.02 | EC | 4.26% |

**Supplementary Table 2. Presence of access path among MtABCG46 WT and mutants**

| Variant | % of presence | Bottleneck radius | |
|---|---|---|---|
| | | Average | StDev |
| WT | 9.08% | 0.97 | 0.06 |
| F562L | 4.49% | 0.98 | 0.07 |
| F562Y | 1.25% | 0.95 | 0.05 |
| F562A | 2.97% | 0.94 | 0.04 |

**Supplementary Table 3. Duration and applied restraints in individual stages of MD simulations**

| MD stage | Simulation time [ns] | Positional restraint energy [Kcal/mol] | | | | Dihedral restraint energy [Kcal/mol] |
|---|---|---|---|---|---|---|
| | | Protein | ATP | $Mg^{2+}$ | Membrane | Membrane |
| Minimization | 5000 steps | 10.0 | 2.5 | 2.5 | 2.5 | 250.0 |
| Equilibration 1 | 1 | 10.0 | 2.5 | 2.5 | 2.5 | 250.0 |
| Equilibration 2 | 1 | 5.0 | 2.5 | 2.5 | 2.5 | 100.0 |
| Equilibration 3 | 1 | 2.5 | 2.5 | 2.5 | 1.0 | 50.0 |
| Equilibration 4 | 1 | 1.0 | 1.0 | 1.0 | 0.5 | 50.0 |
| Equilibration 5 | 1 | 0.5 | 0.5 | 0.5 | 0.1 | 25.0 |
| Equilibration 6 | 1 | 0.1 | 0.5 | 0.5 | - | - |
| Long equilibration | 100 | - | 0.05 | 0.05 | - | - |
| Unrestrained equilibration | 100 | - | - | - | - | - |
| Production | 400 | - | - | - | - | - |

**Supplementary Table 4. MtABCG46 domains defined and residues used to calculate RMSD**

| Region | Residue range | RMSD range |
|---|---|---|
| NBD1 | M1 – L499 | T52 – L478 |
| TMD1 | N500 – L770 | K501 – V769 |
| Linker | G771 - L822 | n/a |
| NBD2 | P823 – P1145 | E825 – V1134 |
| TMD2 | T1146 – R1428 | S1151 – N1424 |
